# Dopamine D1 receptor signalling in dyskinetic Parkinsonian rats revealed by fiber photometry using FRET-based biosensors

**DOI:** 10.1101/2020.03.12.989103

**Authors:** Jace Jones-Tabah, Hanan Mohammad, Shadi Hadj-Youssef, Lucy Kim, Ryan D. Martin, Faïza Benaliouad, Jason C. Tanny, Paul B.S. Clarke, Terence E. Hébert

## Abstract

Like many G protein-coupled receptors (GPCRs), the signalling pathways regulated by the dopamine D1 receptor (D1R) are dynamic, cell-type specific, and can change in response to disease or drug exposures. In striatal neurons, the D1R activates cAMP/protein kinase A (PKA) signalling. However, in Parkinson’s disease (PD), alterations in this pathway lead to activation of extracellular regulated kinases (ERK1/2), contributing to L-DOPA-induced dyskinesia (LID). In order to detect D1R activation *in vivo* and to study the progressive dysregulation of D1R signalling in PD and LID, we developed ratiometric fiber-photometry with Förster resonance energy transfer (FRET) biosensors and optically detected PKA and ERK1/2 signalling in freely moving rats. We show that in Parkinsonian animals, D1R signalling through PKA and ERK1/2 is sensitized, but that following chronic treatment with L-DOPA, these pathways become partially desensitized while concurrently D1R activation leads to greater induction of dyskinesia.

## Introduction

G protein-coupled receptors (GPCRs) play pivotal roles in mediating neuronal communication in the brain. In fact, 90% of non-olfactory GPCRs are found in the brain^1^, where they regulate neuronal activity by engaging a variety of downstream effectors which include second messenger producing enzymes, ion channels, monomeric GTPases and protein kinases. Many GPCRs are pharmacologically targeted in the treatment of neurodegenerative and neuropsychiatric disease, and thus it is critical to understand how these receptors regulate intracellular signalling, and how this in turn regulates circuit function and ultimately, behavior. In cell culture, these signalling pathways have been dissected extensively, through the use of genetically-encoded fluorescent and bioluminescent biosensors. Some of these biosensors have recently been applied *in vivo*, to link specific signalling patterns with behavioral outcomes^2–6^.

In the past decade, the widespread availability of genetically-encoded calcium indicators and complementary *in vivo* imaging technologies has revolutionized the study of neural circuits and behavior. Among these technologies, fiber photometry has emerged as the least invasive, most accessible and affordable approach for recording neural activity in freely moving rodents^7–10^. Many intracellular signalling biosensors have been developed which utilize changes in Förster resonance energy transfer (FRET) between two fluorescent proteins^11^, in order to report levels or activity of second messengers, protein kinases, GTPases, post-translational modifications and protein-protein interactions^12–15^. These tools have been widely used to dissect signaling pathways in cultured cells and have recently begun to be applied *in vivo*, aided by recent technological developments such as intravital imaging^14,16^, microendoscopy^3^, 2-photon microscopy^2,5,6,17^ and fluorescence lifetime measurement^4^. Here we report a ratiometric fiber-photometry approach for real-time recording of genetically-encoded FRET biosensors in freely moving rodents. We apply this approach to investigate alterations in striatal GPCR signalling in a rat model of Parkinson’s disease and L-DOPA induced dyskinesia.

The dopamine D1 receptor (D1R) is a Gα_s/olf_ coupled receptor expressed throughout the forebrain. Like many GPCRs, several factors can impact D1R signalling, including the properties of specific ligands, the cellular context, interactions with signalling partners, and evolving disease processes. The D1R is particularly abundant in the striatum, where it is expressed on medium spiny GABAergic projection neurons that make up the “direct pathway” of the basal ganglia (dMSNs)^18,19^. In dMSNs, D1R activation increases neuronal excitability and promotes synaptic plasticity by stimulating cAMP production and PKA/DARPP-32-dependent phosphorylation of downstream targets. Activation of the direct pathway through D1R thus facilitates movement and promotes synaptic plasticity involved in motor learning^20,21^.

In Parkinson’s disease, degeneration of dopaminergic neurons in the substantia nigra results in striatal dopamine depletion, leading to a hypoactivity of dMSNs. To maintain signalling homeostasis in response to diminished synaptic dopamine levels, dMSNs upregulate expression of D1R-associated effectors including Gα_olf_, effectively sensitizing the receptor and increasing its ability to activate pathways associated with protein kinase A (PKA) and extracellular regulated kinase 1/2 (ERK1/2)^22–25^. Stimulation of such sensitized D1Rs contributes to the development of a common adverse effect of Parkinson’s treatment termed L-DOPA induced dyskinesia (LID)^26–28^. While numerous behavioral and biochemical studies have linked aberrations in D1R signalling to the development of LID, it has not been shown how alterations in protein kinase signalling evolve following degeneration of dopaminergic neurons, or after initiation of dopamine replacement therapy.

To study D1R-dependent signalling in the striatum, we used adeno-associated viral vectors to express genetically-encoded FRET biosensors for PKA and ERK1/2 activity in the dorsal striatum of adult rats and recorded signalling dynamics in behaving animals using a newly developed ratiometric photometry technique. Using a 6-OHDA model of Parkinson’s disease, we show that D1R signalling through these protein kinase pathways is at first enhanced by dopamine depletion, but then attenuated by prolonged L-DOPA treatment. Despite this apparent desensitization of the signalling response, dyskinesia increased in response to D1R stimulation indicating that LID, once established, becomes dissociated from activation of PKA and ERK1/2. Our approach allowed us to monitor signalling in the same animals over the course of dopamine depletion and L-DOPA treatment, providing novel insights into how these signalling processes shape behavioral outcomes over time.

## Results

### Cloning and evaluation of FRET-based protein kinase biosensors AKAR and EKAR in primary rat striatal neurons

We set out to develop a platform for expressing and recording FRET-based biosensors that was both simple to use and readily generalizable to other biosensors and animal models. While transgenic mice have been generated that express a number of FRET biosensors^3,14,29^, we opted to use adeno-associated viral vectors (AAVs) to drive biosensor expression, since this would allow our tools to be exploited in both rats and mice without the need to maintain colonies of transgenic animals. In this work, we used previously published FRET-based reporters for PKA and ERK1/2 referred to as AKAR and EKAR respectively^13^. Variants of these biosensors have previously been used *in vivo* with microendoscopy^3,13^, 2-photon microscopy^5^, and fluorescence lifetime photometry^4^. These biosensors are substrates for protein kinase activity and contain phosphorylation-dependent interaction domains such that the biosensor undergoes a conformational change upon phosphorylation, leading to an increase in resonance-energy transfer (Fig. 1A). We subcloned AKAR and EKAR constructs into an AAV backbone containing a neuron-specific promoter and used AAV serotype 1 to transduce primary striatal neurons from rat brain (Fig. 1B-E). We confirmed neuron-specific expression by immunofluorescent labelling (Supplemental Fig. 1), and we also performed live-cell FRET imaging in 96 well plates using high-content confocal microscopy (Fig. 1B-E). In primary neurons, the AKAR-associated FRET ratio was increased by the addition of the D1R agonist SKF 81297 or the adenylyl cyclase activator, forskolin (Fig. 1C). The PKA response to SKF 81297 but not forskolin was inhibited by pre-treatment with the D1R antagonist SCH 23390, but not the MEK inhibitor U0126. Conversely, the EKAR-associated FRET ratio was increased by addition of NMDA or forskolin, and was attenuated by pre-treatment with the MEK inhibitor U0126 (Fig. 1D). This pattern of results is consistent with previous studies indicating that D1R couples only weakly to ERK1/2 under normal physiological conditions^24^. Finally, the elevated PKA activity seen after acute SKF 81297 administration could be partially reversed by subsequent addition of a D1 antagonist, demonstrating that the FRET response can be reversed when signalling is blocked (Fig. 1E).

**Figure 1:**
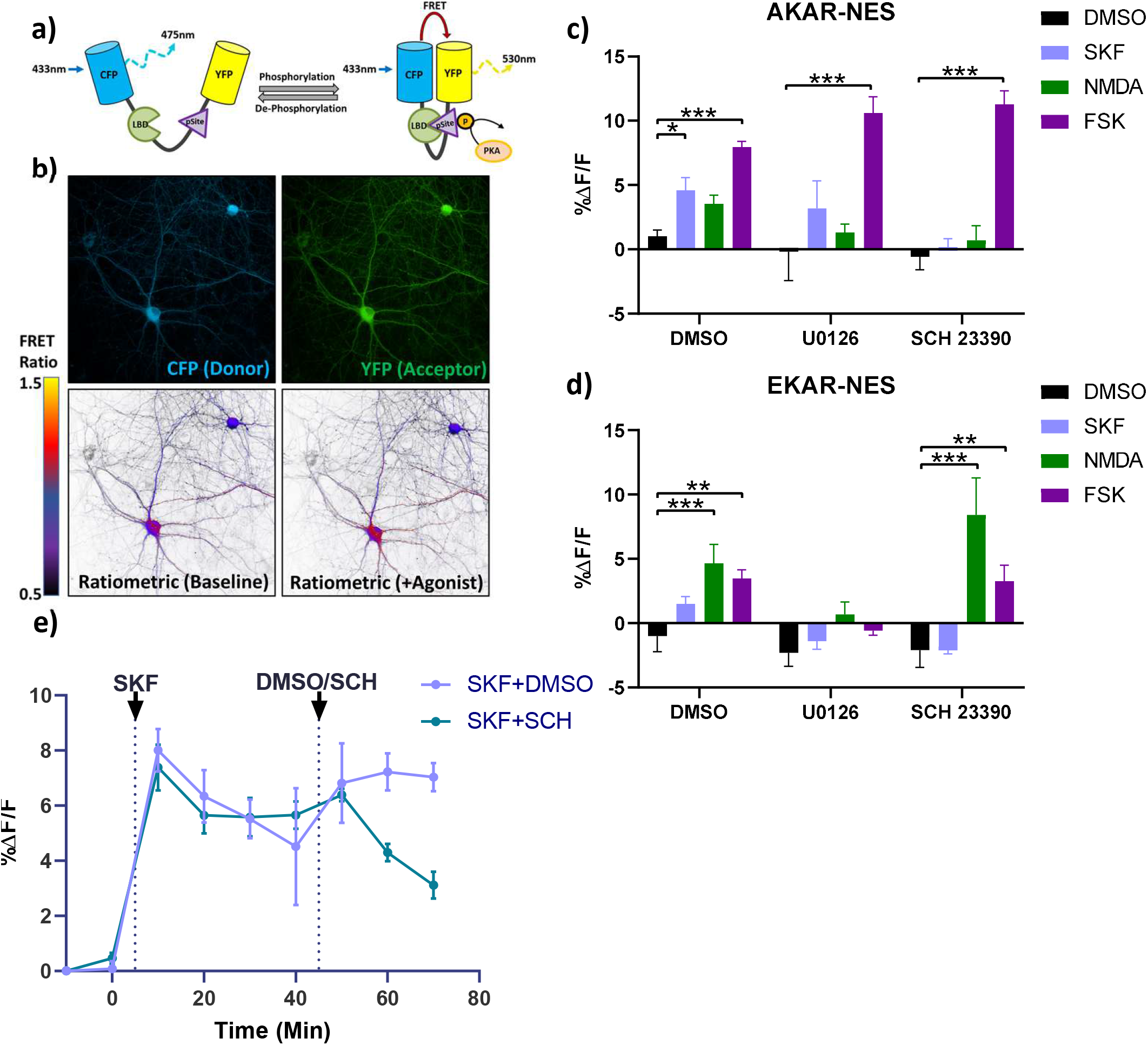
Evaluation of FRET-based protein kinase sensors AKAR and EKAR for detecting signalling pathway activation in primary neuronal cultures. **(a)** Schematic depiction of the PKA biosensor, showing how the molecular rearrangement that results from PKA phosphorylation in turn leads to changes in FRET signal. The ERK1/2 biosensor EKAR works by an analogous mechanism. **(b)** Representative images showing neurons transduced with AKAR. These images show the CFP donor (top left), YFP acceptor (top right), and ratiometric pseudo-images before and after treatment with the D1R agonist SKF 81297 (lower left and right, respectively). Ratiometric images were generated by plotting the calculated FRET ratio (YFP/CFP) at each image pixel according to the color scale shown. **(c, d)** Bar graphs depicting change in PKA (c) or ERK1/2 (d) signalling in primary cultured neurons following a 5-min treatment with the indicated drugs and displayed as change in FRET relative to baseline FRET (%ΔF/F). **(e)** Time course of activation of PKA in response to D1R agonist SKF 81297, followed by reversal with D1R antagonist SCH. Imaging was performed at approximately 10-min intervals due to the time required to image all wells on the multiwell plate. Abbreviations and drug doses: DMSO, dimethyl-sulfoxide, SKF, SKF 81297 (1 μM), NMDA, N-methyl-D-aspartic acid (5 μM), FSK = forskolin (5 μM), U0126 (1 μM), SCH = SCH 23390 (1 μM). Data are displayed as mean ± SEM for 3-6 independent experiments. *p < 0.05, **p < 0.01, ***p < 0.001 vs. DMSO within each pre-treatment group (Bonferroni-corrected t-tests).

### *In vivo* expression of FRET biosensors in rat striatum

To express FRET biosensors *in vivo*, we injected AAV-AKAR (titer ~5×10^12^ viral genomes/ml) into the dorsal striatum of adult Sprague-Dawley rats. We then assessed expression 3 weeks later by immunofluorescence microscopy in fixed brain sections (Fig. 2A-B). Fluorescently labelled cell bodies stained positive for NeuN and DARPP-32, but not the astrocytic marker GFAP, suggesting that expression of the biosensor was restricted to neurons (Fig. 2B, Supplemental Fig. 2A-C). In the rat striatum, 95% of neurons are medium spiny GABAergic projection neurons that express DARPP-32^30,31^. Of these, approximately half express D1R and contribute to the direct pathway projecting from the striatum to the substantia nigra reticulata (SNr), whereas the remainder express D2R and make up the indirect pathway projecting to the external globus pallidus (GPe)^32,33^. In a sagittal brain section, we observed fluorescently labelled axons projecting to both the GPe and SNr, suggesting transduction of neurons giving rise to either pathway (Supplemental Fig. 2D).

**Figure 2:**
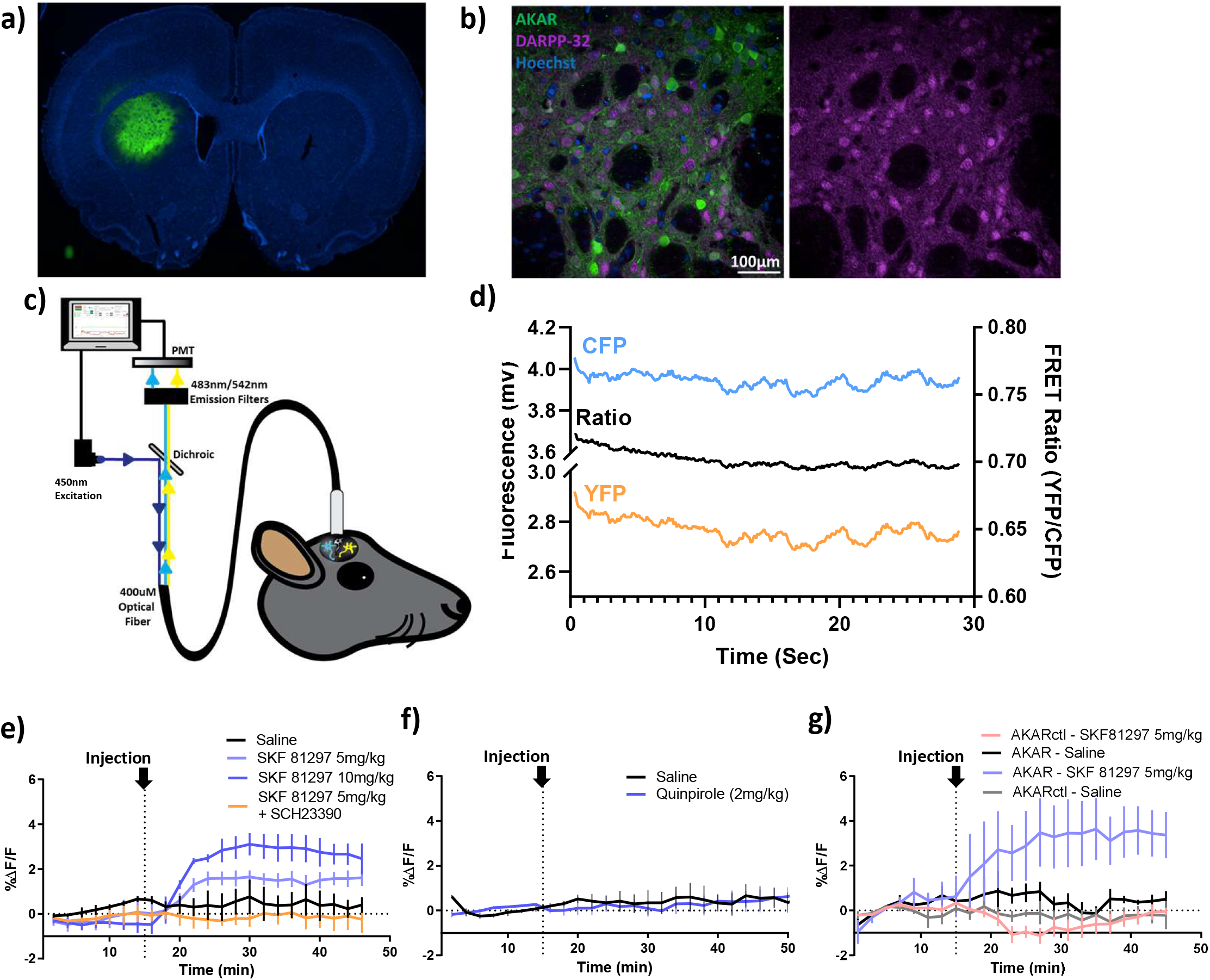
AAV-mediated expression of AKAR in rat striatum and photometry recording in anesthetized and freely moving rats. **(a)** Representative image showing expression of AKAR in a coronal section containing the dorsal striatum (i.e. caudate-putamen) of an adult rat that was sacrificed 3 weeks following stereotaxic injection of AAV-AKAR. Imaging was performed on fixed tissue sections, with AKAR fluorescence shown in green and cell nuclei stained with Hoechst shown in blue. **(b)** Representative higher-magnification image showing fluorescent DARPP-32 immunolabelling of fixed striatal tissue, together with AKAR expression. **(c)** Schematic representation of the ratiometric photometry system. **(d)** Exemplar trace showing CFP and YFP channels, and calculated FRET ratio, recorded for 30 s from AKAR-transduced rat dorsal striatum at 100 Hz sampling rate. The lower panels show photometry recordings obtained in rats maintained either under light isoflurane anesthesia **(e, f)** or freely moving **(g)**, with FRET values averaged within consecutive 30-s time bins. **(e)** After 15 min of recording, rats were injected s.c. (vertical dotted line) with vehicle or the D1R agonist SKF 81297, with or without 30-min pre-treatment with the D1R antagonist SCH 23390. **(f)** Rats were injected with vehicle or the dopamine D2R agonist quinpirole. Responses were recorded for 30 s every 2 min, for at least 30 min post-injection. **(g)** Photometry recording performed in freely-moving rats with either AKAR or AKARctl, the latter being a negative control biosensor lacking a phosphorylation site. Rats were recorded at 2-min intervals, and were challenged with either vehicle or SKF 81297 (5 mg/kg s.c.), 15 min after the start of recording. For all experiments, individual rats were recorded under vehicle and drug conditions on consecutive days in a counterbalanced design. Panels **e-g** show the mean ± SEM (n=3 rats), where the value for each rat is the mean FRET over a 30 s long 100-Hz recording such as is shown in panel D. Refer to *Methods* for full description of data processing and representation.

### Development of CFP/YFP ratiometric photometry

*In vivo* imaging of FRET biosensors has previously been accomplished using microendoscopy^3,14^, 2-photon microscopy^5^ and fluorescence lifetime photometry^4^, but these techniques are costly, technically demanding and require specialized equipment that make them impractical for wide application in non-specialized labs. In contrast, fiber photometry is a simple method for performing functional fluorescence recording *in vivo* and is compatible with many common behavioral tests^7–9^. We therefore worked with Labeo Technologies (Montréal) to develop a fiber-photometry system for performing ratiometric FRET measurements through surgically implanted optical cannulae (schematically shown in Fig. 2C). Excitation and emission filters were selected based on the experimentally determined emission spectra of AKAR expressed in HEK 293 cells (Supplemental Fig. 3). Using excitation with a 450 nm laser-diode, we detected fluorescence signals corresponding to the FRET donor (CFP) and acceptor (YFP) from AAV-AKAR injected rats that were >3-fold greater than background, which was recorded from control rats which had a probe implanted but were not injected with virus (Supplemental Fig. 4). An example of the raw fluorescence trace and corresponding FRET ratio acquired at 100 Hz from a single rat expressing the AKAR sensor is shown in Fig. 2D. To minimize the potential for photobleaching during long duration recordings, laser illumination was applied in discrete 30-s intervals followed by an interval with the laser turned off (Supplemental Fig. 5A). The mean fluorescence from each of these 30-s recording intervals was then averaged in order to condense the data for analysis. Furthermore, illumination of brain tissue at 450 nm was found to produce approximately 3-fold greater autofluorescence in the CFP channel compared to the YFP channel, and this autofluorescence bleached upon extended illumination (Supplemental Fig. 5B). To remove this background autofluorescence, we recorded from two age-matched sham-injected control rats in each experiment and subtracted the average background signal corresponding to the time of recording for each FRET experiment (further described in *Materials and Methods* and shown schematically in Supplemental Fig. 5C).

**Figure 3:**
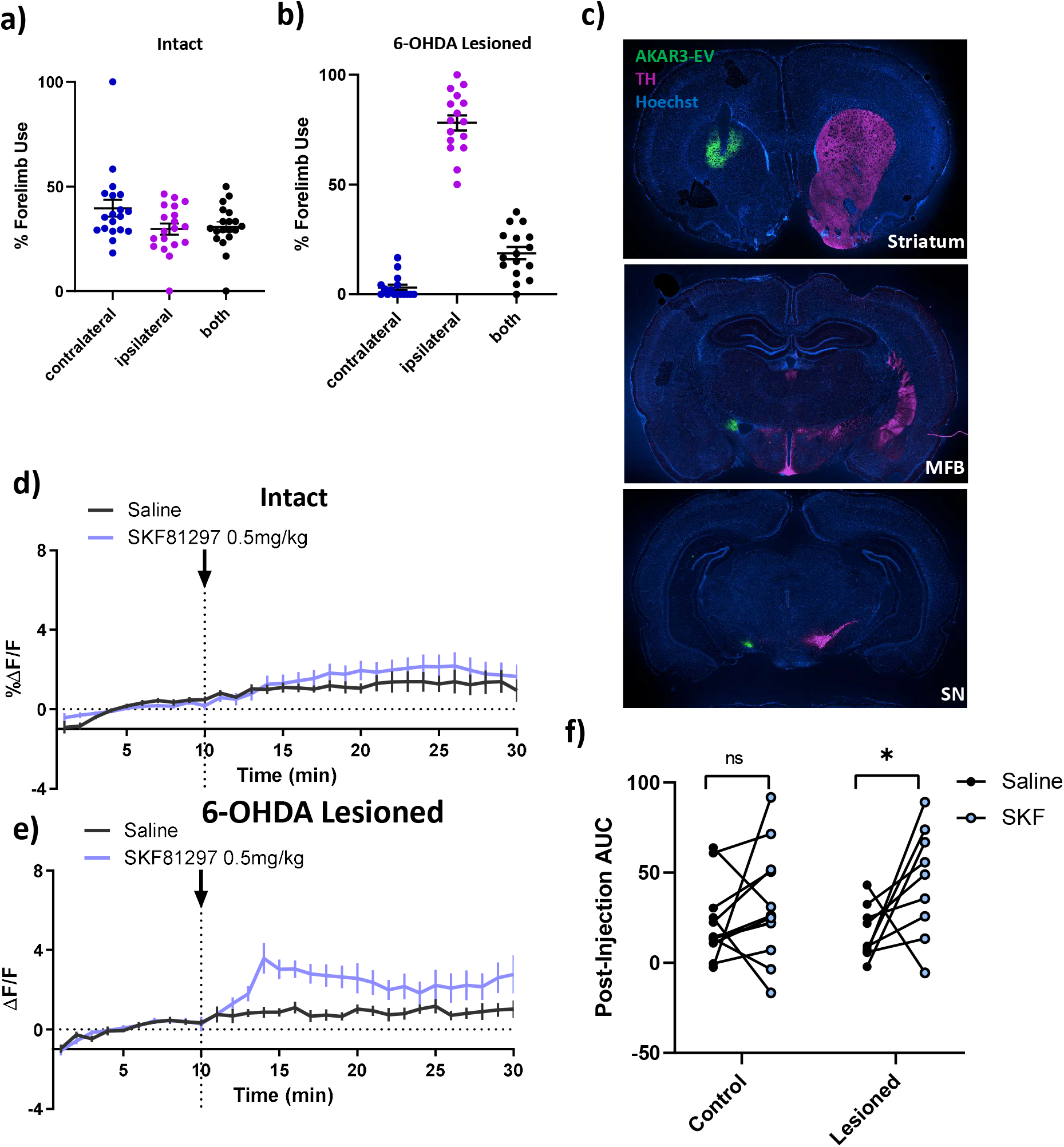
Using photometry to measure D1R-dependent PKA activation in intact and 6-OHDA lesioned hemi-Parkinsonian rats. Panels **a** and **b** show cylinder test results for intact rats and 6-OHDA lesioned rats, respectively. Rats in both groups had been implanted with optic probes and expressed AKAR in the dorsal striatum, in the case of 6-OHDA injected animals this was ipsilateral to the lesion. Y-axes show the percentage of rears in which individual animals used the forelimb that was ipsilateral *vs*. contralateral to the AAV-injected hemisphere, or used both forelimbs at the same time. Tests in the 6-OHDA lesioned rats occurred 3 weeks following intra-striatal injection of AAV_1_-SynTet-AKAR and optical probe implantation. **(c)** Representative immunofluorescent images depicting AKAR expression (green) and optic-probe placement in the dorsal striatum in addition to expression of tyrosine hydroxylase (TH, shown in purple) in the striatum, medial forebrain bundle (MFB) and substantia nigra (SN). **(d)** FRET recordings performed on intact rats expressing AKAR and injected with either saline or the D1R agonist SKF 81297 (0.5 mg/kg s.c.) at t = 10 min (n = 12). **(e)** FRET recordings performed on 6-OHDA lesioned rats expressing AKAR and injected with either saline or SKF81297 (0.5 mg/kg s.c.) at t = 10 min (n = 9). For all experiments, rats were recorded under both saline and drug conditions in a counterbalanced design. **(f)** Area under the curve (AUC) analysis of panels d and e. AUC was calculated relative to baseline following drug or saline injection. AUC values for Saline *vs*. SKF 81297 in each control and lesioned rats were by repeated measures t-tests, * p < 0.05.

### D1R agonist dose-dependently activates PKA signalling in dorsal striatum

To test the ability of this *in vivo* system to detect agonist-induced effects on protein kinase signalling, we expressed AAV-AKAR in the dorsal striatum and recorded responses to systemically-administered D1 and D2 agonists (Fig. 2E-G). Photometric recording was initially performed with rats under light anesthesia in order to immobilize the animals and isolate drug-induced changes from behavior-associated fluctuations in striatal signaling that might occur in moving animals. Recording in these anesthetized rats, we observed a rapid, and dose-dependent increase in PKA activation following subcutaneous injection of the D1R agonist SKF 81297 (Fig. 2E). In contrast, quinpirole, an agonist for D2 receptors, which are coupled to Gα_i_ rather than Gα_s_, elicited no response (Fig. 2F). The lack of negative FRET response to D2R activation in anesthetized rats is consistent with prior studies showing that PKA signalling in the indirect pathway is basally maintained at low levels and is not further reduced by D2R activation^34^. We next tested freely moving rats with SKF 81297 and found similar, albeit more variable, increases in the FRET ratio (Fig. 2G). In order to confirm that the observed changes in FRET were due to specific signalling effects, we expressed a control AKAR biosensor in which the phosphorylation substrate threonine residue had been mutated to an alanine residue. No agonist-induced effect was observed in freely moving rats that expressed the control biosensor (Fig 2G).

### D1R-dependent PKA activation is enhanced in a 6-OHDA lesion model of Parkinson’s disease

The histopathological hallmark of Parkinson’s disease (PD) is degeneration of nigral dopaminergic neurons, which in turn causes the main clinical symptoms of rigidity, tremor and akinesia^35^. Patients with PD do not typically present with motor deficits until they have lost at least 50% of dopamine neurons, and an even greater number of striatal dopamine terminals^36^. One potential explanation for this phenomenon is that compensation takes place through increased sensitivity of post-synaptic dopamine receptor signalling. In support of this hypothesis, lesion of the nigrostriatal dopamine pathway in animal models increases the expression of postsynaptic D1R effectors such as Gα_olf_^22,23,25^, accompanied by behavioral and biochemical sensitization to D1R agonists^24,37^.

We therefore sought to determine whether we could observe enhanced D1R-dependent PKA signalling in a rat model of PD. To this end, we used 6-hydroxydopamine (6-OHDA) to unilaterally lesion midbrain dopamine neurons^38^. A hemi-Parkinsonian phenotype was confirmed, first behaviorally using the cylinder test of forelimb use asymmetry (Fig. 3A-B), and subsequently by tyrosine hydroxylase immunostaining (Fig. 3C). In a separate group of rats, we also verified that expression of AAV-AKAR, together with optic probe implantation in the striatum, did not itself alter performance in the cylinder test (Fig 3A). We expressed AAV-AKAR in the striatum of the lesioned hemisphere and recordings were performed 4 weeks post-lesion. Rats were injected s.c. with SKF 81297, at a dose (0.5 mg/kg s.c.) that was previously shown to induce striatal cFos expression in 6-OHDA lesioned but not intact rats^24^. Accordingly, we observed SKF 81297-induced PKA activation only in 6-OHDA lesioned rats (Fig. 3D-F), indicating that D1R-mediated cAMP signalling was enhanced following a period of dopamine depletion.

### D1R dependent PKA and ERK1/2 activation is enhanced by dopamine depletion and desensitized following chronic L-DOPA

The gold standard of treatment for motor impairment in PD patients remains the dopamine precursor L-DOPA. However, long-term treatment with L-DOPA frequently leads to the emergence of acute L-DOPA-induced dyskinesia (LID), a debilitating adverse drug effect characterized by abnormal involuntary movements^39,40^. Dyskinesia can be driven acutely by optogenetic activation of dMSNs in 6-OHDA lesioned rodents^41–43^ and the development of LID is dependent on D1R activation^26^. In particular, in animal models LID has been found to depend on sensitized D1R signalling through PKA and ERK1/2^27,28^, but it remains unclear whether the primary role of D1R-dependent protein kinase activity in LID development is direct, by phosphorylating synaptic targets and acutely increasing excitability, or indirect, through long term D1R-driven changes in synaptic plasticity and intrinsic excitability of dMSNs.

To investigate the role of D1R-dependent PKA and ERK1/2 activity in LID, we measured D1R signal pathway engagement in dopamine-depleted and drug naïve animals, and then compared this to the activation of the same pathways following two weeks of “priming” with daily L-DOPA injections which has previously been shown to produce dyskinesia in 6-OHDA lesioned rats^44,45^. We therefore expressed AKAR or EKAR biosensors in the striatum of hemi-parkinsonian rats, and started testing 4 weeks post-lesion, as described in the previous section (see Fig. 4A for the complete timeline). In drug naïve 6-OHDA lesioned rats, we recorded significant activation of both PKA and ERK1/2 in response to SKF 81297 (0.5 mg/kg s.c.) but not saline injection (Fig. 4B,C). Rats were then treated daily for two weeks with L-DOPA (6 mg/kg s.c.) and the DOPA decarboxylase inhibitor benserazide (15 mg/kg s.c.) to induce dyskinesia. After 2 weeks of L-DOPA priming, rats displayed a sensitized acute dyskinetic response to a subsequent L-DOPA injection (Fig. 4D). Acute dyskinesia was measured at 10-min intervals using the previously described abnormal involuntary movements (AIMS) scoring system^46^, and was found to peak approximately 60 min after injection, and return to baseline 180 min after injection (Fig. 4D); this finding is consistent with previous studies using this model^44,45^. During this acute dyskinesia, PKA activity also increased and then returned to baseline at 180 min post-injection. We next re-tested the response to SKF 81297 in these L-DOPA primed rats after a 24 or 48 hour abstinence from L-DOPA. In comparison to drug naïve animals, D1R agonist-induced PKA activation was attenuated at both time points (Fig. 4E).

**Figure 4:**
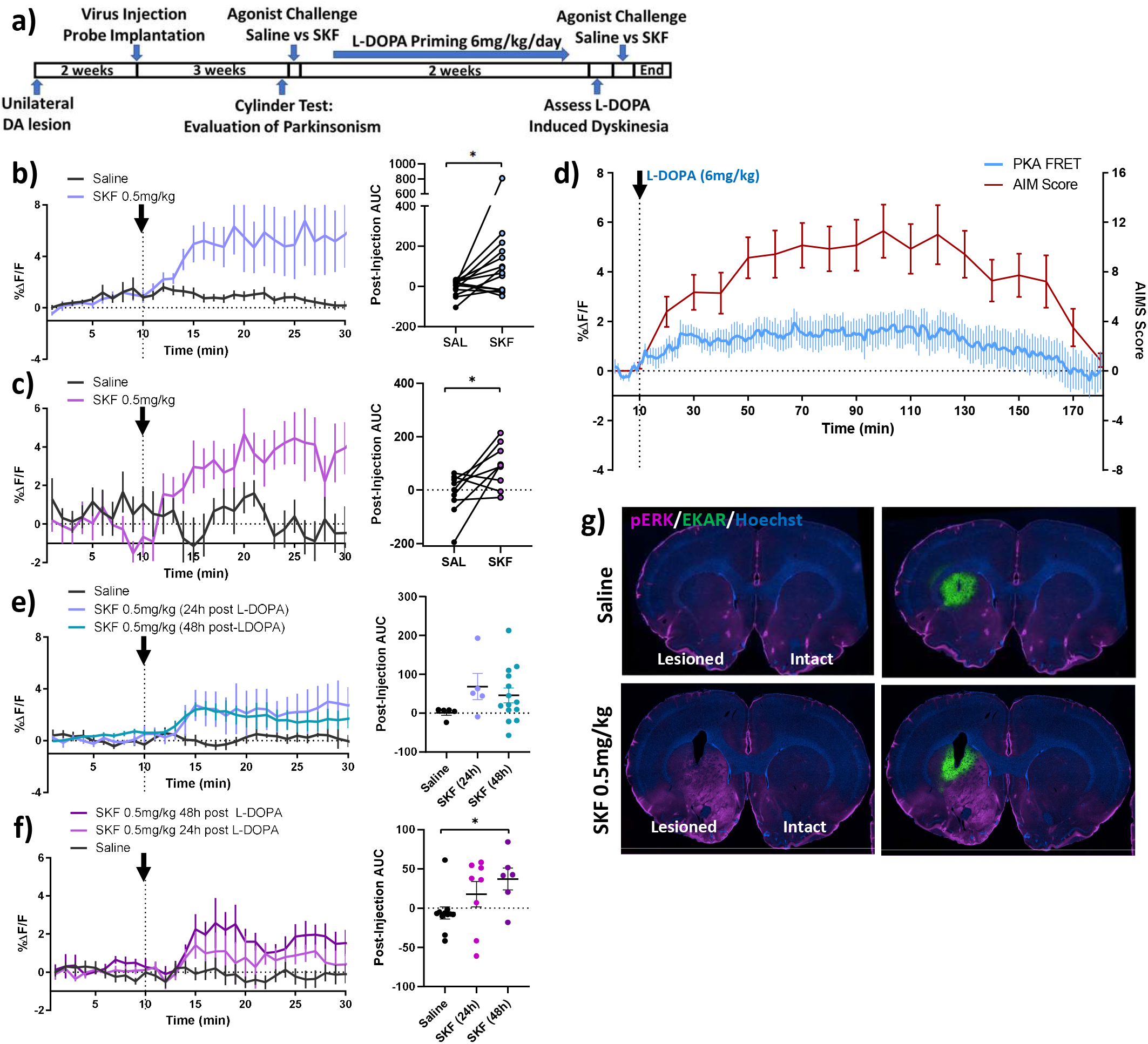
PKA and ERK1/2 signalling in Parkinsonian rats and following development of L-DOPA-induced dyskinesia. **(a)** Timeline of surgeries, drug treatments and testing for L-DOPA-induced dyskinesia (LID) experiments. **(b)** PKA biosensor responses to saline or SKF 81297 (0.5 mg/kg s.c) in drug naive 6-OHDA lesioned rats. The left panel shows time-course of FRET responses, while the right panel shows the integrated area-under the curve (AUC) for the post-injection period. All animals were tested under both saline and drug conditions, as indicated by the connecting lines on the AUC plot (n = 14 rats). **(c)** ERK1/2 biosensor responses to saline or SKF 81297 in drug naive 6-OHDA lesioned rats. The left panel shows the time-course of FRET responses, while the right panel shows the integrated area-under the curve (AUC) for the post-injection period. All animals were tested under both saline and drug conditions, as indicated by the connecting lines on the AUC plot (n = 8 rats). **(d)** 6-OHDA lesioned rats develop L-DOPA-induced dyskinesia following 2 weeks of daily treatment with L-DOPA/benserazide (6/15 mg/kg s.c.). Dyskinesia was measured as an AIM score every 10 min (red line), while PKA activation was simultaneously recorded at 2 min intervals (blue line) (n = 14 rats). **(e)** PKA biosensor responses to saline or SKF 81297 in L-DOPA primed, dyskinetic rats, either 24 or 48 hours after the last L-DOPA treatment. The left panel shows time-course of FRET responses, while the right panel shows the integrated area-under the curve (AUC) for the post-injection period (n = 5, 14). **(f)** ERK1/2 biosensor responses to saline or SKF 81297 in L-DOPA primed, dyskinetic rats, either 24 or 48 hours after the last L-DOPA treatment. The left panel shows time-course of FRET responses, while the right panel shows the integrated area-under the curve (AUC) for the post-injection period (n = 6-8 rats). **(g)** Representative immunofluorescent images depicting phospho-ERK1/2 (pERK) immunolabelling in L-DOPA primed, dyskinetic rats challenged with either saline (n = 3) or SKF 81297 (n = 4), 24 hours following the last L-DOPA treatment. Data in all time-course plots are mean ± SEM. Statistical analysis in panels b and c was performed by paired t-test. Statistical analysis in panels e and f was performed by paired t-test with Bonferroni correction, * p < 0.05.

For ERK1/2, we found that activation was attenuated 24 hours after L-DOPA but was partially restored 48 h after L-DOPA (Fig. 4F). We additionally sought to validate our signalling data by immunolabelling for phosphorylated (active) ERK1/2 (Fig. 4G). Following the last photometry recording experiment, rats continued to receive daily L-DOPA for 1 week. Twenty-four hours after the last L-DOPA injection, rats were injected with either saline or SKF 81297 and perfused 30 min after injection. In saline injected rats, there was no detectable difference in phospho-ERK labelling between the lesioned and intact striata. However, SKF 81297 in fact did induce phospho-ERK selectively in the lesioned hemisphere, an effect that was not detected in our photometry experiment. This could indicate a difference between measuring phosphorylation of ERK itself versus measuring ERK protein kinase activity relative to phosphatase activity. However, it also suggests that while our technique offers the advantage of tracking changes in signalling in the same animals over time, there is a threshold below which effects may be undetectable by photometry.

### Repeated L-DOPA enhances dyskinesia but attenuates PKA and ERK1/2 activation induced by D1R activation

Concurrent with FRET recordings, we compared the behavioral effects of D1R activation in the drug naïve and L-DOPA treated animals. In drug naïve rats, SKF 81297 induced moderate dyskinesia within 30 min of injection, whereas activation of PKA occurred as early as 5 min post-injection (Fig. 5A). In contrast, after 2 weeks of L-DOPA treatment, the SKF 81297-induced dyskinesia had doubled in magnitude, while activation of PKA was suppressed (Fig. 5B). The same trend was also observed for ERK1/2 activation (Fig. 5C-D). In order to perform statistical comparisons, we calculated the integrated area-under the curve for the signalling data and confirmed that while PKA and ERK1/2 activation were significantly decreased by L-DOPA treatment, dyskinesia (measured as the integrated AIM score 30 min after drug injection) was significantly increased (Fig. 5E-H).

**Figure 5:**
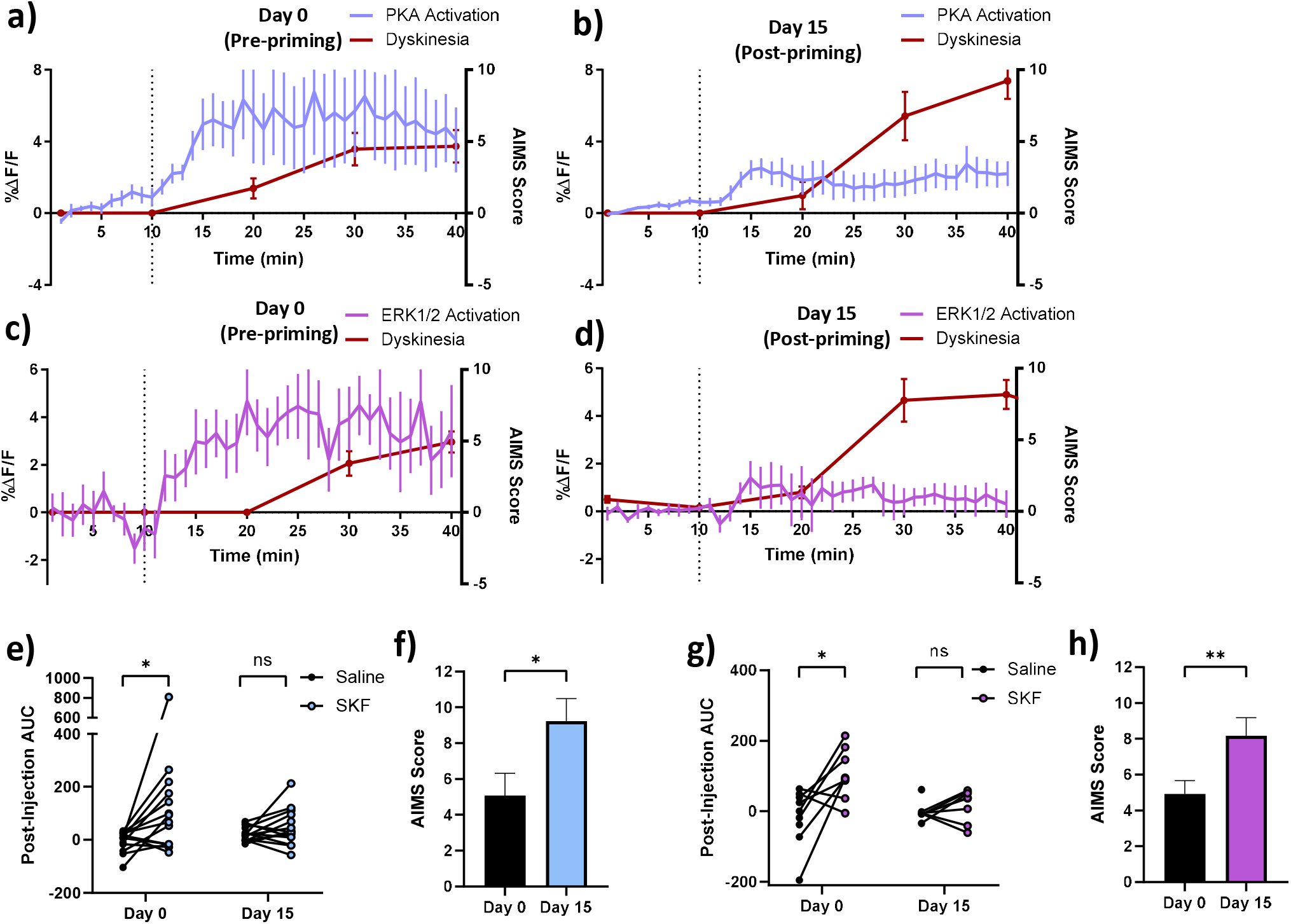
D1R agonist-induced PKA and ERK1/2 signalling is significantly attenuated following chronic L-DOPA, while dyskinesia is significantly enhanced. PKA and ERK1/2 biosensor response profiles (%ΔF/F) and dyskinesia scores (AIMS) in drug naïve, 6-OHDA lesioned rats and L-DOPA primed, dyskinetic rats treated with the D1R agonist SKF 81297 (0.5 mg/kg s.c.) **(a)** PKA biosensor response profiles and dyskinesia scores in drug-naïve, 6-OHDA lesioned rats treated with SKF 81297 (0.5 mg/kg s.c.) (n = 14 rats) **(b)** PKA biosensor response profiles and dyskinesia scores in L-DOPA primed, dyskinetic rats treated with SKF 81297 (n = 14 rats). **(c)** ERK1/2 biosensor response profiles and dyskinesia scores in drug naïve, 6-OHDA lesioned rats treated with SKF 81297 (n = 9 rats) **(d)** ERK1/2 response profile and dyskinesia score in L-DOPA primed, dyskinetic rats treated with 0.5mg/kg SKF81297 (n = 8 rats). **(e)** Integrated area-under the curve (AUC) for the post-injection period comparing PKA activation by saline versus SKF 81297 on Day 0 (pre-priming) and Day 15 (post-priming). **(f)** Comparison of peak dyskinesia (measured at 40 min post-injection) on Day 0 versus Day 15 in AKAR rats. **(g)** Integrated area-under the curve (AUC) for the post-injection period comparing ERK1/2 activation by saline versus SKF 81297 on Day 0 (pre-priming) and Day 15 (post-priming). **(h)** Comparison of peak dyskinesia (measured at 40 min post-injection) on Day 0 versus Day 15 in EKAR rats. All data is presented as mean ± SEM. Statistical analysis of photometry data before and after L-DOPA priming (e, g) was performed using paired t-test with Bonferroni correction. For EKAR data, one rat was excluded from the repeated measures ANOVA due to a missing value. Comparison of dyskinesia (f, h) was performed by paired t-test, * p < 0.05, **p<0.01.

At the completion of the study, all animals were perfused and brain sections were analyzed by immunofluorescence with tyrosine hydroxylase staining to ensure the correct anatomical location of the virus and optical probe and confirm the loss of striatal dopamine terminals (data not shown). We additionally found that hallmark biochemical markers of L-DOPA induced dyskinesia, FosB and Ser-10 phosphorylated histone 3, were increased in the lesioned striatum following chronic L-DOPA^47–49^, further validating our LID model (Supplemental Fig. 6).

## Discussion

To our knowledge, our study is the first to employ ratiometric fiber photometry to study GPCR signalling in live animals. More specifically, we provide a proof-of-concept application of *in vivo* monitoring of GPCR-dependent protein kinase activity, using ratiometric fiber photometry in combination with FRET-based biosensors of PKA and ERK1/2 activity. This approach revealed D1R-dependent activation of these signalling pathways in freely moving rats. Concordant with previous work, we observed increased activation of PKA and ERK1/2 by D1R in the striatum of 6-OHDA lesioned rats. A chronic regimen of L-DOPA/benserazide that induced dyskinesia also resulted in attenuation of this enhanced signalling, even though D1R-associated dyskinesia was significantly increased. The occurrence of D1R supersensitivity is well established in 6-OHDA lesioned rats [reviewed in^50,51^], and a recent study using FRET-based biosensors has demonstrated hyperactivation of PKA and ERK1/2 by D1R agonists in striatal slices taken from 6-OHDA lesioned mice^52^. The present *in vivo* study extends these *ex vivo* findings, by showing that acute D1R-induced PKA and ERK1/2 activation is suppressed by repeated L-DOPA treatment, suggesting that while D1R signalling may be desensitized by chronic L-DOPA treatment, the D1R-driven dyskinesia nevertheless increases.

### Comparison with other approaches

In the past two decades, genetically-encoded FRET biosensors have served as a critical tool in revealing spatially and temporally dynamic signalling networks regulated by GPCRs in cultured cells. Many FRET-based sensors have now been created, allowing the optical detection of a wide range of signalling processes including second messengers, ion flux, protein kinases, monomeric GTPases, post-translational modifications and protein-protein interactions^12–15,53^. In contrast, the *in vivo* application of intracellular signalling-related FRET biosensors is much less developed, especially in freely-moving animals.

Various technologies have been developed in recent years to enable *in vivo* fluorescence imaging in the brain. Efforts to date have tended to focus on measuring calcium and voltage dynamics with single-fluorophore, intensity-based biosensors. However, FRET-based biosensors are also compatible with a range of imaging technologies, including 2-photon intravital imaging, micro-endoscopy with fibered or head mounted microscopes, and fiber photometry. For example, intravital microscopy, which allows sub-cellular resolution imaging of live cells in native tissue, can be used in transgenic mice expressing FRET biosensors to image signalling dynamics in cell populations throughout the body and brain^2,14,16,54,55^. However, for imaging neurons in conscious animals, the subject needs to be head-fixed to the microscope, an invasive procedure that greatly limits the behavioral repertoire that can be tested. Furthermore, 2-photon imaging approaches are limited to superficial layers of the cortex unless tissue is resected, or a lens is implanted within the brain to reach deeper structures. Epi-fluorescent imaging can be performed on freely moving animals using an endoscope consisting of an implanted lens connected via optical fiber to a conventional microscope, or a head mounted mini-scope. The former approach was previously used to image PKA and ERK1/2 signalling in the nucleus accumbens using transgenic mice expressing the same biosensors used in our study ^3^.

The principal advantages of the ratiometric fiber photometry method reported here relate to its simplicity and ease of use. Compared to imaging approaches, photometry occupies a distinct technical niche by sacrificing spatial resolution in favor of reduced cost and invasiveness, a greater range of concommitant behavioral readouts, and the potential for higher throughput or multiplexed recordings from multiple brain regions, which can be accomplished by adding channels to the device^56,57^. Ratiometric photometry requires minimal expertise in optics or image processing, and can be combined in principle with any FRET biosensor in order to study a range of intracellular signalling processes. Furthermore, by using a fiberoptic rotary joint to allow free rotation of the animal being recorded, this technique is highly compatible with rotational behaviors that are a feature of the unilateral 6-OHDA lesion model employed in our study.

### Challenges

Potential difficulties associated with recording FRET signals *in vivo* include possible tissue autofluorescence and the differential scattering of the donor and acceptor wavelengths of light as they traverse biological tissue. In our hands, the main drawback of this technique was its limited ability to detect small changes in signalling. As described in Fig. 4G we found that an increase in phospho-ERK that was detectable by immunofluorescence was not reported by photometry recording with the EKAR sensor. This could represent a limitation in the sensitivity of the assay, or it could reflect differences in measuring abundance of phospho-ERK by immunofluorescence (where signals are amplified) versus measuring net ERK1/2 protein kinase activity by FRET, which takes into account opposing phosphatase activity. Limitations in sensitivity can be addressed by designing more sensitive biosensors with a larger dynamic range. Indeed, a more sensitive AKAR biosensor was recently developed by adding a subcellular localization signal to target the biosensor to PKA-rich locations within the cell^5^. Another limitation of our study was that biosensor expression, although neuron-specific, was not restricted to the neuronal population responsive to D1 agonists, i.e. D1R-expressing neurons. Since the latter represent less than 50% of all striatal neurons, our signal-to-noise ratio could likely be improved through more targeted expression, for example using an appropriate line of Cre-driver transgenic animals.

An additional challenge is the potential for hemodynamic artifacts to affect biosensor outputs. Notably, in a small number of animals, there was a significant *reduction* in fluorescence intensity detected in both CFP and YFP channels after administration of the D1R agonist. Given that D1R agonists can impact blood flow^58–60^ we hypothesized that this effect was caused by alterations in local hemodynamics around the optical probe. Interference from local hemodynamics in fluorescent imaging has been previously reported both with FRET-based and single fluorophore, intensity-based biosensors, and experimental and computational approaches have been proposed to address this confounding factor^53,61,62^. In our experiments, the decrease in fluorescence signal caused by the presumed hemodynamic changes led to a reduction in FRET ratios, which could mask the effects of D1R activation. Future studies will be required to explore solutions to each of these challenges.

FRET can be measured both ratiometrically, as in our study, and as a function of fluorescence lifetime. The latter approach requires recording only from the donor of a FRET pair, and because the lifetime of a fluorophore is an intrinsic property of the molecule, lifetime-based measurements are more quantitative compared to ratiometric FRET^63^. Measuring FRET by fluorescence lifetime could circumvent some of the challenges described above. Photometry systems based on time-correlated single photon counting have previously been described^7^ and it was recently shown that this approach could be used to measure PKA activity in the nucleus accumbens using a FRET biosensor optimized for lifetime-based detection^4,64^. Like the imaging techniques described previously, the main drawback of lifetime approaches are the increased equipment costs and complexity of the technique.

### Mechanisms underlying L-DOPA induced dyskinesia

L-DOPA-induced dyskinesia is a complex phenomenon involving multiple neurotransmitters and signalling pathways. Nevertheless, alterations in D1R signalling appears to play a central role and the development of LID is causally dependent on D1R activation and signalling^26,41–43^. In animal models based on dopamine depletion, D1R signalling through Gα_olf_ becomes more responsive to agonist stimulation, resulting in increased activation of PKA. This sensitization is associated with altered D1R surface expression and trafficking, increased expression of Gα_olf_ with a loss of negative regulatory mechanisms^22,24,37,65–68^.

Previous studies using pharmacological approaches have demonstrated the importance of PKA and ERK1/2 in the development of dyskinesia. For example, in 6-OHDA lesioned rats, intra-striatal inhibition of PKA during L-DOPA treatment was found to attenuate both LID and its molecular correlates such as ΔFosB accumulation, p-DARPP-32 and pERK1/2^27^. Similar results have also been shown with inhibitors of ERK1/2 phosphorylation given during chronic L-DOPA treatment^28,69^. Interestingly, while these studies all found that inhibition of PKA or ERK1/2 during the L-DOPA priming period reduced LID, inhibition of ERK1/2 on the day of dyskinesia testing was not required to attenuate LID^28^. Taken together, these data suggest that the primary role of exaggerated PKA and ERK1/2 signalling is in the development, rather than the expression, of LID.

Our findings here suggest that while D1R coupling to PKA and ERK1/2 is increased in dopamine depleted rats, the ability of D1R to activate these pathways is attenuated following two weeks of L-DOPA treatment. This desensitization of PKA and ERK1/2 activation nevertheless co-occurs with increased coupling of D1R activation to the induction of dyskinesia, indicating that once LID has developed there is a dissociation between the activation of these signalling pathways by D1R and the behavioral response.

The increased coupling of D1R activation to LID despite desensitization of these signalling pathways may reflect PKA- and ERK1/2-dependent remodelling of the direct pathway at the level of gene expression and synaptic function. Our *in vivo* observations support a model in which activation of PKA and ERK1/2 during initial exposures to L-DOPA prime dMSNs to become hyper-reactive to subsequent stimulation. This model is supported by electrophysiological studies which have shown that treatment with L-DOPA or D1R agonists normally reverses the hypoactivity of dMSNs associated with dopamine depletion, whereas the development of dyskinesia leads to hyperactivity of these neurons above normal levels^43,70^. This persistent hyperactivity of dMSNs in LID is thought to be the result of an inability to reverse corticostriatal long term potentiation (LTP), which in turn is regulated by D1R and PKA^71–73^. In addition to regulating synaptic function, D1R-dependent activation of PKA and ERK1/2 also regulates gene expression. One of the hallmarks of dyskinesia is accumulation of the AP-1 transcription factor subunit ΔFosB, which appears to have a causal role in LID^47,74,75^. In fact sustained overexpression of ΔFosB in the striatum of Parkinsonian primates was found to replicate behavioral and molecular markers of LID in the absence of chronic L-DOPA treatment^76^. These data support a model in which hyperactivation of PKA and ERK1/2 in the dopamine depleted striatum lead to persistent dyskinesia, despite the fact that these pathways are themselves desensitized in dyskinetic animals.

## Conclusion

We have presented a novel method of ratiometric fiber photometry for monitoring intracellular signalling in freely moving animals using FRET-based biosensors. The system described here is simple to construct and use, and could be paired in principle with any FRET biosensor. We have applied this approach to measure sensitization of D1R-induced PKA and ERK1/2 signalling in the striatum following a period of dopamine depletion. The technique further allowed us to demonstrate that L-DOPA treatment partially reverses the sensitization of these signalling pathways, but nevertheless results in greater coupling of D1R activation to the induction of acute dyskinetic symptoms.

## Materials and Methods

### Animals

Adult male Sprague-Dawley rats and Sprague-Dawley dams with postnatal day 0 pups were purchased from Charles River, Saint-Constant QC, Canada. Animals were maintained on a 12/12-hour light/dark cycle and free access to food and water. All procedures were approved by the McGill University Animal Care Committee, in accordance with Canadian Council on Animal Care Guidelines.

### Drugs

For all *in vitro* experiments, SKF 81297 HBr (Toronto Research Chemicals), N-methyl-D-aspartate (NMDA, Sigma-Aldrich), forskolin (Sigma-Aldrich), SCH 23390 HCl (Tocris) and U0126 (Tocris), were initially dissolved in DMSO to prepare stock solutions and then diluted to the desired concentration in artificial cerebrospinal fluid. For *in vivo* experiments, SKF 81297 HBr, quinpirole HCl and SCH 23390 HCl (Tocris) were dissolved in physiological saline and injected in a volume of 1 ml/kg. For 6-OHDA lesion experiments, desipramine HCl (Sigma-Aldrich), and pargyline (Sigma-Aldrich) were dissolved in physiological saline and injected at a volume of 1 ml/kg. 6-OHDA HBr (MilliporeSigma) was prepared at 7 μg/μL dissolved in 0.02% ascorbic acid and 0.9% NaCl, and stored at −80°C for up to one week prior to use. For dyskinesia experiments, L-DOPA methyl ester HCL and benserazide (Sigma) were dissolved in 0.9% saline with 0.1% ascorbate and injected s.c. at 1 ml/kg.

### Cloning and Virus Production

The FRET-based protein kinase sensors AKAR3EV-NES/NLS and EKAREV-NES/NLS were generously provided by Dr. Michiyuki Matsuda^13^. The pAAV-SynTetOFF plasmid used for all adeno-associated viral constructs was a generous gift from Dr. Hiroyuki Hioki^77^. Subcloning was performed by insertion of a custom multicloning site into the pAAV-SynTetOFF plasmid followed by restriction digestion subcloning to insert each of the biosensors into the AAV backbone.

For cell culture experiments and preliminary *in vivo* validation (Figs. 1 and 2), adeno-associated virus production, purification and concentration was performed as previously described^78^. Briefly, HEK 293T cells were cultured in DMEM containing 10% fetal bovine serum and 1% penicillin/streptomycin until they were 80% confluent. Calcium phosphate was used to co-transfect cells with pAAV transfer plasmid, pXX680 adenoviral helper plasmid and pXR1 packaging plasmid in a 1:3:1 molar ratio. Cells were harvested 60 hours post-transfection and lysed by 3 cycles of freeze-thaw in AAV lysis buffer. Crude AAV preparation was purified by ultracentrifugation on an OptiPrep™ density gradient and concentrated in PBS using Amicon® Ultra centrifugal filters. To determine the AAV titer, viral DNA was extracted using a phenol-chloroform extraction and quantified by qPCR using primers targeted at the viral WPRE sequence. For a detailed protocol, refer to^78^. For experiments involving 6-OHDA lesioned animals (Figs. 3-5), all viruses were produced by the Neurophotonics Platform Viral Vector Core at Laval University.

### Primary Neuronal Culture and AAV Transduction

Primary striatal neurons were prepared from freshly dissected Sprague-Dawley rat pups at postnatal day 0. The day before dissection, 96-well optical bottom imaging plates (Nunc) were coated with 0.1 mg/ml poly-D-lysine overnight. All solutions were prepared and stored at 4°C. Prior to dissection, plates were washed three times with sterile water and allowed to dry, and solutions were warmed to 37°C. The rat pups were decapitated on ice, and brains were rapidly removed and placed in ice cold Hank’s balanced salt solution (HBSS) without calcium and magnesium (Wisent). Striata were dissected and digested for 18 min while rotating at 37°C with papain (Sigma) diluted to a final concentration of 20 units/ml in a neuronal medium (Hibernate A Minus Calcium, BrainBits). Tissue was then transferred to HBSS containing 10% fetal bovine serum, 12 mM MgSO_4_, and 10 units/ml DNase1 (Roche) for 2 min to halt digestion, and then triturated in HBSS with 12 mM MgSO_4_, and 10 units/ml DNase1 using a fire-polished Pasteur pipette until a cell suspension was produced. The cell suspension was passed through a 40 μm mesh filter (Fisher) to remove undigested tissue and then centrifuged on an OptiPrep™ gradient as previously described^79^ to remove cell debris and non-neuronal cells such as microglia. Purified neurons were then diluted in Neurobasal™-A Medium (NBA) with 1× final concentration of B27 supplement (Gibco), 1% GlutaMAX (Gibco) and 1% penicillin/streptomycin (henceforth referred to as complete NBA) supplemented with 10% fetal bovine serum, and 50,000 cells were plated in a total volume of 75 μl per well. Sixteen hours after plating, cells were washed with warm HBSS and media was changed for complete NBA containing 5 μM cytosine-D-arabinoside to inhibit glial cell growth. Cultures were maintained in complete NBA, and media was refreshed by exchanging 30% of the volume with fresh media every 3 days. Primary striatal neurons were transduced by adding AAV directly into the culture media three days after cell plating, using a multiplicity of infection of 5000 viral genomes/cell. Neurons were then cultured for 7-10 days prior to imaging.

### FRET Imaging of Primary Cultures

One hour prior to imaging, media was exchanged for artificial cerebrospinal fluid (ACSF) as previously described ^80^ and cells were returned to the incubator. Imaging was performed at 37°C and 3% CO_2_ using an Opera Phenix™ high-content confocal microscopy system (Perkin Elmer). Images were acquired at 40x magnification using a 425 nm laser for excitation of CFP and emissions detected at 435-515 nm (CFP) and 500-550 nm (YFP). All drugs were initially dissolved in DMSO and then diluted 1:1000 in ACSF. After baseline images were acquired, drug treatment was performed by adding the diluted drugs 1:10 to achieve the final concentration in the well. All image analysis was performed using Columbus™ analysis software (Perkin Elmer). Briefly, transduced cells were automatically identified based on CFP fluorescence intensity, and then filtered based on morphological properties to exclude non-cell objects. CFP and YFP intensities were determined for each pixel and used to calculate the FRET ratio (YFP/CFP) at each pixel and then the mean FRET for each cell. Between 100-2000 transduced neurons were analyzed per well and used to calculate the mean FRET ratio per well at each time-point. For each well, changes in FRET were expressed as ΔFRET (baseline – stimulated) divided by baseline FRET, and expressed as a percentage (%ΔF/F). Each experiment was performed with at least 2 technical replicates (i.e. 2 wells on the plate receiving the same treatments).

### Stereotaxic Surgery

Male Sprague-Dawley rats (Charles River) weighing 275-300 g were anesthetized with isoflurane (3% induction, 1% maintenance) mixed with oxygen, mounted in a stereotaxic apparatus and maintained under isoflurane for the duration of surgery. A midline scalp incision was made and the skull was cleaned in order to visualize bregma and lambda. The angle of the head was adjusted so that the skull was flat and bregma and lambda were at the same height. Using a hand-held micro drill (NeuroStar) with 0.6 mm bit, a hole was drilled for viral injection and probe placement in the dorsal striatum. Additional skull holes were drilled for the insertion of 5 stainless steel surgical screws. Adeno-associated virus (titer ~5×10^12^ viral genomes/ml) was injected at a volume of 1-2 μl and a rate of 0.2 μl/min using a Hamilton syringe and 26-gauge needle, with the tip located at the following coordinates relative to bregma: +1.2 AP, ±2.7 ML and −5.5 DV. For this purpose, a NeuroStar microinjection robot was mounted to the stereotaxic frame. After injection, the syringe needle was left in place for 10 min and then slowly removed. An optic probe (Doric Lens MFC_400/430-0.48_5mm_MF2.5_FLT, 0.4 mm external diameter) was then implanted, with the tip positioned 0.5 mm dorsal to the virus injection site. Once the optical probe was in place, dental cement was used to secure it to the surgical screws. Rats received postoperative monitoring and were administered carprofen analgesic (10 mg/kg s.c.) during surgery and every 24 hours for 3 days after surgery. Subsequent testing was performed a minimum of 2-6 weeks post-surgery, to allow expression of the biosensor.

For 6-OHDA lesion experiments, stereotaxic surgery was performed in the same manner, except as follows. Thirty minutes prior to surgery, rats received a subcutaneous injection of pargyline (5 mg/kg) in order to inhibit 6-OHDA degradation, and desipramine HCl (10 mg/kg) to inhibit 6-OHDA uptake into noradrenergic terminals^81^. An injection of 6-OHDA was made in the right medial forebrain bundle (MFB) at the following coordinates relative to bregma: −2.8 AP, 2.0 ML, −9.0 DV. Each rat received 2.5 μl of 6-OHDA HBr (7 μg/μL in 0.02% ascorbic acid dissolved in 0.9% NaCl; MilliporeSigma). 6-OHDA was injected at a rate of 0.5 μL/min, with the needle withdrawn after a 10-min delay. The wound was closed with surgical sutures. Animals were allowed to recover for two weeks before behavioral evaluation of Parkinsonism.

### Assessment of Parkinsonism

Parkinsonian signs were assessed in the cylinder test, as previously described^81^. Rats were placed individually in a transparent glass cylinder (14 cm diameter × 28 cm height), and observed for 5 min. During this time, an observer noted the number of weight-bearing wall contacts made with each forelimb during rearing, and a video was recorded for later behavioral analysis. Only animals exhibiting preferential use of the ipsilateral forepaw (i.e. in ≥ 70% of rears) were selected to undergo the AAV stereotaxic injections within one week of forelimb testing followed by dyskinesia induction three weeks after AAV injection.

### Histology and immunofluorescence

Rats were perfused with 0.1% phosphate-buffered saline (PBS) for 10 min followed by 4% paraformaldehyde solution in PBS, pH 7.4, for 10 more min. The brains were extracted and maintained in 4% paraformaldehyde solution for 24 h at 4°C and then transferred to 0.1% PBS at 4°C. Coronal or parasagittal sections (30 μm thick), which included the striatum or substantia nigra, were collected using a vibratome (TPI series 1000) at room temperature (RT). Free-floating sections were first washed three times in PBS and then permeabilized for 10 min with 0.3% Triton-100 in PBS. Antigen retrieval was performed by transferring the sections to 80°C citrate buffer for 30 min, then allowing them to cool to RT. Sections were washed once with PBS and then blocked overnight at 4°C with 10% bovine serum albumin (BSA) in PBS. Primary antibodies were incubated overnight at 4°C in PBS 10% BSA. Alexa-fluor conjugated secondary antibodies (Invitrogen) were incubated for 3 hours at RT in PBS containing 5% BSA. Nuclei were stained for 10 min at RT with Hoechst 3334 (H3570, Thermo Fisher Scientific) diluted 1:10,000 in PBS, and sections were washed twice more in PBS. Sections were then mounted on microscope slides using Fluoromount mounting medium (E126367, Invitrogen) and stored at 4°C until imaged. Mounted sections were imaged on the Opera Phenix™ High Content microscope (Perkin Elmer) using either 5× or 40× objectives. Images acquired at 5× were stitched using the Harmony software (Perkin Elmer) to generate whole brain section images.

The details of the antibody dilutions and sources are as follows. Primary antibodies were: anti-GFP (1:500; sc-8334) and anti-fos-B (1:1500; sc-48), both from Santa-Cruz; anti-histone H3 pS10 (pH3) (1:300; ab47297), from Abcam; anti-tyrosine hydroxylase (1:1000; P40101), from Pel-Freez. Anti-DARPP-32 (1:1000, #2302), from Cell Signalling; anti-GFAP (1:1000; GA524), from Agilent. Secondary antibodies were: anti-mouse Alexa 488 (1:500; A21236) and anti-rabbit Alexa 647 (1;500; A21245), both from Invitrogen.

### Construction of ratiometric photometry recording system

The FRET acquisition system was designed and built by Labeo Technologies (Montréal, QC, Canada). The excitation source is a laser diode centered at 450 nm with an optical filter at 438 ± 12 nm. A dichroic mirror (with a cut-off wavelength of 458 nm) allows separation of excitation and emission channels. On the detection side, two photomultiplier tubes (PMTs, Hamamatsu Photonics) are used to measure fluorescence. Each detection channel is coupled with an emission filter (483 ± 16 nm for CFP and 547 ± 27 nm for YFP) and another dichroic mirror (cut-off wavelength of 520 nm) is used to split them. PMT signals are amplified with trans-impedance amplifiers and acquired with a data acquisition device (National Instruments) at up to 1 kHz, with the possibility to record up to 4 auxiliary analog inputs in synchrony. The acquisition electronics are connected to a computer through a USB port. Both excitation and emission optical pathways are co-aligned and connected to a 400 μm optical fiber which is connected to a 400 μm fiberoptic patch cable (Doric Lens), which can be connected to the animal directly, or with an intervening rotary joint (Doric Lens).

The acquisition user interface is used to control laser power, acquisition frequency and program the duration and frequency for periodic recordings. A Matlab analysis script provided with the FRET acquisition system is used to open saved acquisitions for post-processing and allows visualization of the recordings and generation of basic descriptive statistics. Following processing in Matlab, the data is exported in CSV format for subsequent analysis.

### Photometry recording and analysis

For recordings in anesthetized animals, rats were rapidly anesthetized in an induction chamber using 3% isoflurane and then maintained at 0.5% isoflurane while mounted loosely on a stereotaxic frame with a nose-cone to supply oxygen and isoflurane. Under this level of anesthesia rats were unconscious but maintained normal reflexive responses. The photometry system patch cable was connected to the surgically implanted optic probe using a ceramic mating sleeve (Doric Lens). Baseline recordings of at least 10 min were acquired prior to drug administration. For experiments in freely moving rats, a fiberoptic rotary joint (Doric) was mounted above the test chamber allowing the rats to freely rotate while connected to the photometry patch cable.

All photometry recording was performed using laser power of 20 μW and with PMT gain set to 70% for both the CFP and YFP channels. To minimize bleaching or photo-toxicity, the laser was turned on for only 30 out of every 60 s (for test sessions <60 min long) or 30 out of every 120 s (for longer test sessions). Data was initially processed using a custom MatLab script to stitch together individual 30-s recordings and downsample the data from 1000 Hz to 100 Hz for subsequent analysis.

Data from each AKAR and EKAR recording were analyzed using a custom R script we wrote to perform the following series of operations. The first step was to subtract background autofluorescence which was found to decrease over time in response to sustained illumination. Accordingly, age-matched control (sham-injected) rats were used to calculate background CFP and YFP values; these control rats were recorded under identical conditions to experimental subjects. The mean background for CFP and YFP was plotted as a function of time (Supplemental Fig. 5B) and fit to a linear equation which was then used to subtract CFP and YFP background at each time-point. After background subtraction, the FRET ratio was calculated as CFP/YFP. Baseline FRET was calculated as the mean FRET for the pre-injection period and all measurements were subsequently expressed as the percentage ΔFRET relative to baseline FRET [i.e. %ΔF/F = 100*(FRET − FRET_baseline_)/ FRET_baseline_]. The mean %ΔF/F for each experimental group were then plotted. Area under the curve was calculated from the %ΔF/F *vs*. time plots using the following formula, where *C* is the %ΔF/F value corresponding to each timepoint (*T*), and B is defined as the mean %ΔF/F of the baseline period.

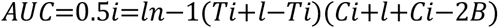

### Drug Treatments during photometry recording

Drug treatments of saline versus agonist in photometry experiments were generally performed in a counterbalanced, repeated measures design with each animal being tested under both saline and drug conditions on consecutive days, with the following exception. In Fig. 1e, the SKF 81297 (10 mg/kg) and SCH 23390 tests were performed on separate days in a non-counterbalanced fashion. Antagonist pre-treatments were administered 30 min prior to the beginning of a test session. All drug injections during photometric recordings were performed during a laser-OFF interval and without disconnecting the animal from the recording system. Experimenters were not blinded to drug conditions.

### Dyskinesia induction and AIM assessment

In order to model L-DOPA induced dyskinesia, Parkinsonian rats were primed to elicit abnormal involuntary movements (AIMs) with once-daily injections of L-DOPA/benserazide (6/15 mg/kg s.c.) for 14 days. AIMs were then assessed after the last injection (Day 14). To this end, rats were placed in glass cylinders and AIMs were scored for 2 min at 10-min intervals. L-DOPA/benserazide (6/15 mg/kg) was injected 10 min after rats were placed in the cylinder and each test lasted for a total of 3 h. Orolingual, limb and axial dyskinesias were scored according to the scales as previously described ^46^. The integrated AIMs score was then calculated as the sum of the intensity score multiplied by the duration score for each of the behaviors scored (orolingual, limb and axial). The same procedure was used to score AIMS in rats treated with Saline or SKF 81297 (0.5mg/kg). Simultaneous photometry recording was performed during all dyskinesia experiments.

### Statistical Analysis

All statistical testing was performed using Prism 8 (GraphPad). For cell culture experiments, data for each biosensor was analyzed first by 2-way ANOVA, with multiple comparisons performed using Bonferroni-corrected t-tests. Area-under-the-curve analysis of *in vivo* recording data was performed as follows. For comparison of saline *vs*. SKF 81297 at single time-points (Fig. 4b, 4c), data were analyzed by paired t-tests. For comparisons of saline *vs*. SKF 81297 at multiple time-points before and after L-DOPA (Fig. 4e, 4f, 5e, 5g), multiple comparisons were made by Bonferroni corrected paired t-tests comparing each SKF 81297 condition to saline. Comparison of dyskinesia score before and after L-DOPA treatment was performed by paired t-test (Fig. 5F, 5H).

## Acknowledgements

This work was supported by grants from the Weston Brain Institute and the NSERC ENGAGE program J.J.T was supported by doctoral studentships from the CIHR and the McGill Healthy Brains for Healthy Lives initiative. R.M. was supported by studentships from the McGill-CIHR Drug Development Training Program and the McGill Faculty of Medicine. S.H.Y. was supported by a summer scholarship from NSERC. P.B.S.C. is a member of the Center for Studies in Behavioral Neurobiology at Concordia University, Montreal. We thank Dr. Philippe Huot (McGill Neurological Institute) for guidance in implementing the 6-OHDA rat model, Dr. Michiyuki Matsuda (Kyoto University) for providing us with biosensor constructs and Dr. Hiroyuki Hioki (Kyoto University) for providing AAV plasmids. Lastly, we thank members of the Hébert, Clarke and Tanny labs for feedback and guidance throughout the development of the project, and critical reading of the manuscript.

## Author Contributions

JJT, PBSC and TEH designed the experiments and wrote the manuscript. JJT, HM, SHY, LK and FB conducted experiments. RDM, HM and JJT analyzed experiments. JJT, HM, PBSC, RDM, JCT and THE edited the manuscript.

## Competing Interests

None

---

**Figure S1:**
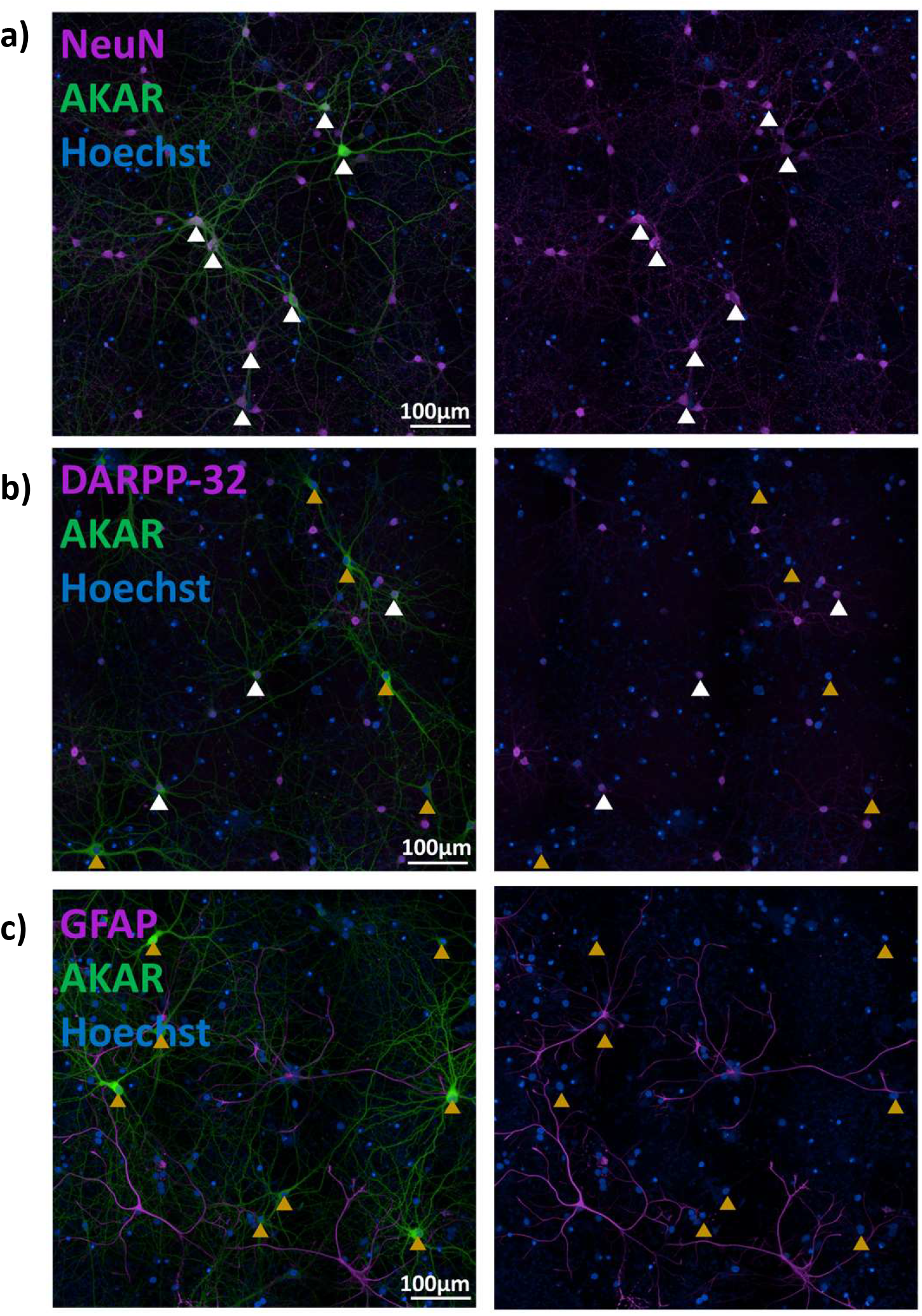
AAV_1_-SynTet-AKAR expresses in NeuN and DARPP-32 positive cells, but not in GFAP positive cells in primary neuronal culture. Representative images showing primary neurons transduced with AAV_1_-SynTet-AKAR (green) and immunolabelled with either a general neuronal marker, NeuN (panel A), a marker of striatal medium-spiny GABAergic neurons, DARPP-32 (panel B), or an astrocytic marker, GFAP (panel C). White and orange triangles respectively indicate AKAR expressing cells that are positively or negatively labelled for the indicated proteins. Nuclei are labelled with Hoechst stain (blue). Representative images were selected from the set of images acquired by high-content imaging of primary neurons plated on 96-well plates. At least 3 wells were stained for each of NeuN, DARPP-32 and GFAP, and more than 25 images per well were acquired.

**Figure S2:**
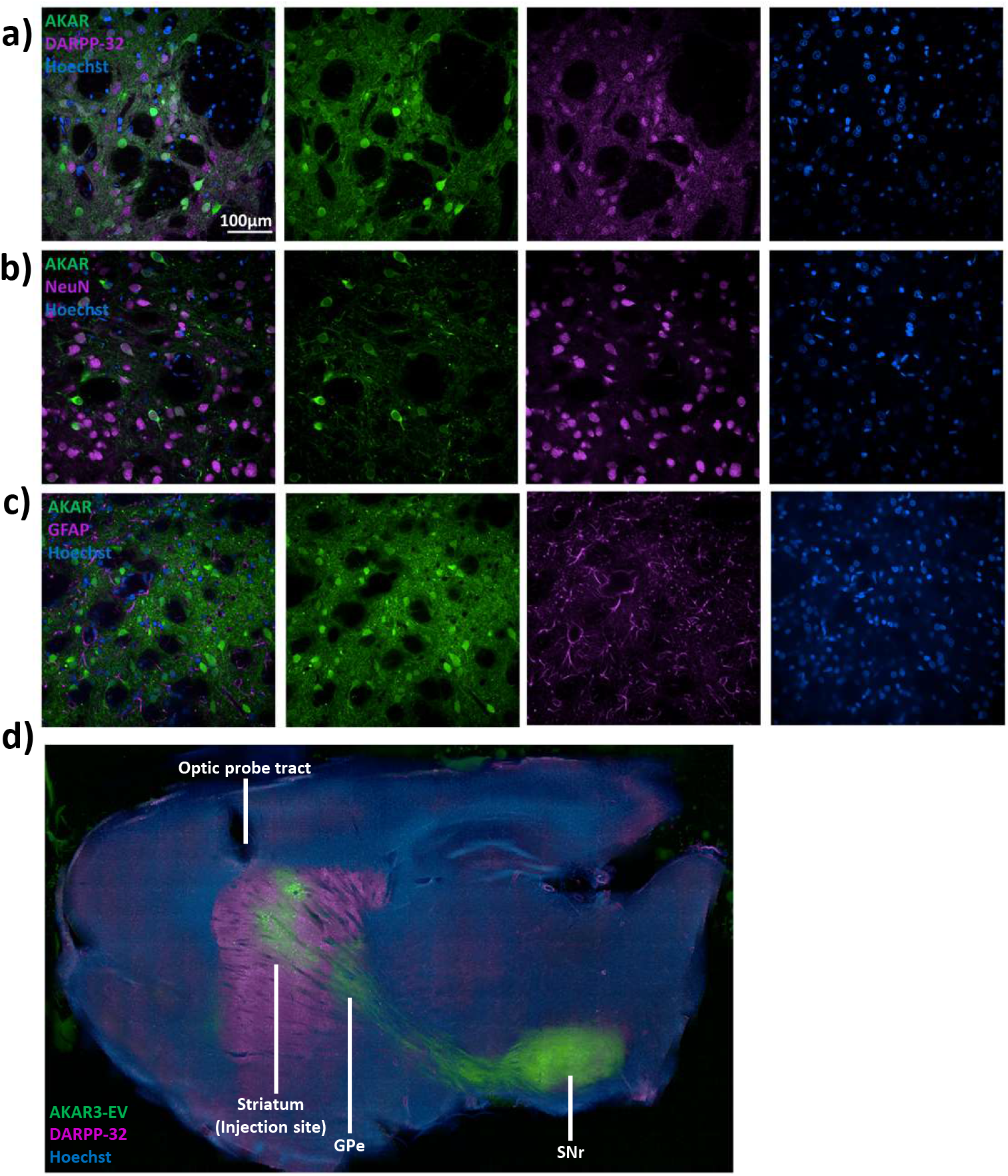
AAV_1_-SynTet-AKAR expresses in NeuN and DARPP-32 positive, but not GFAP positive cells in the dorsal striatum. **(a-c)** Representative images of fixed striatal tissue sections from rats injected with AAV_1_-SynTet-AKAR (green) and immunolabelled with a general neuronal marker, NeuN (a), a marker of striatal medium-spiny GABAergic neurons, DARPP-32 (B), or an astrocyte marker, GFAP (c). Nuclei are labelled with Hoechst stain (blue). **(d)** Sagittal section from a rat injected in the dorsal striatum (caudate/putamen) with AAV_1_-SynTet-AKAR (green). White lines indicate fluorescently labelled axons projecting from the striatum to the target nuclei of the indirect pathway and direct pathways, the external globus pallidus (GPe) and substantia nigra reticulata (SNr) respectively.

**Figure S3:**
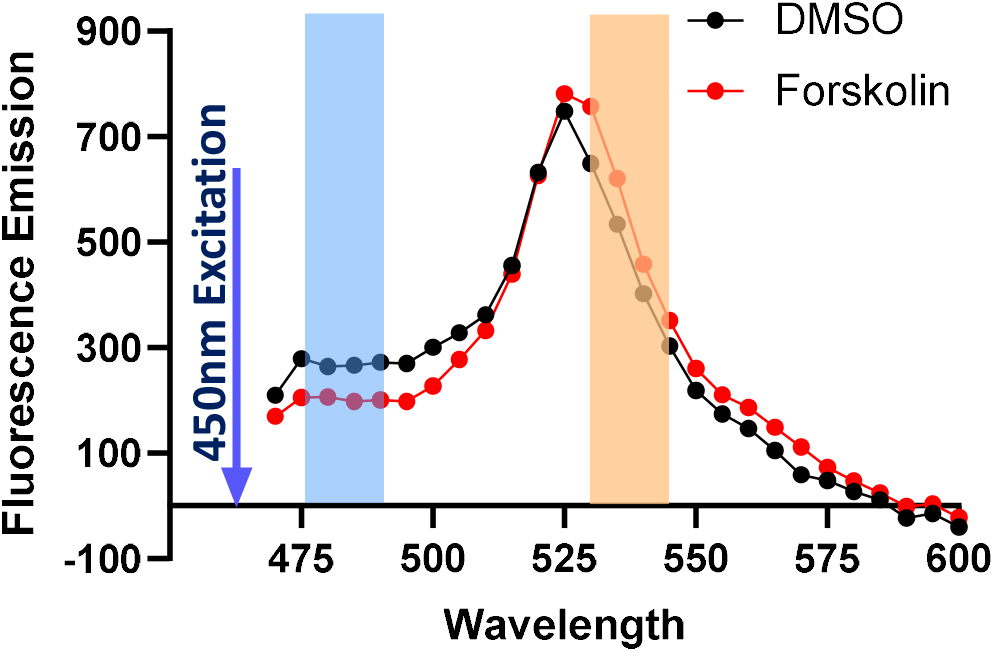
FRET sensor emission spectra. Emission spectra of AKAR expressed in HEK 293 cells under 450 nm excitation. Emission spectra were determined under basal conditions (DMSO vehicle) and following treatment with a saturating concentration of the adenylyl cyclase activator forskolin. Areas shaded in blue and orange represent the spectral range of the emission filters used for CFP and YFP in the photometry system (see *Methods* for additional details). Fluorescence emission is shown in arbitrary units.

**Figure S4:**
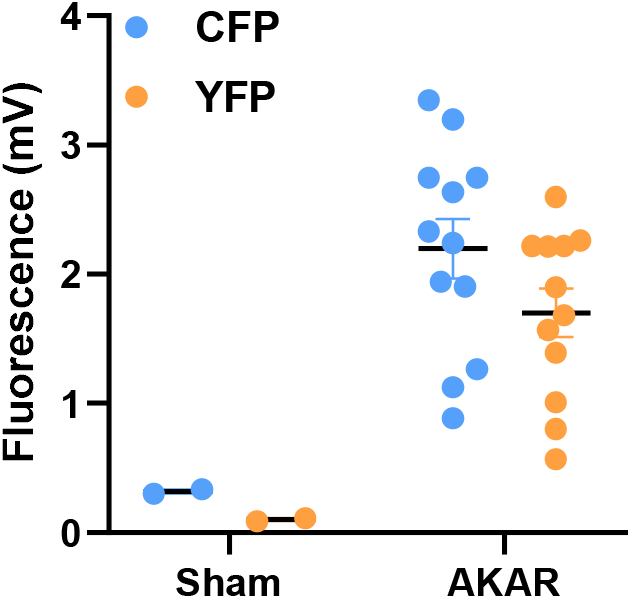
Detection of fluorescence *in vivo* in probe-only *vs*. AKAR-injected rats. Intensity of emissions recorded by photometry in the CFP and YFP channels using 450 nm excitation at 20 μW laser power. Recordings were made in either rats with optical probes only or 3 weeks after injection of AAV_1_-SynTet-AKAR. Fluorescence intensity is measured as photomultiplier tube output and is displayed as mV units.

**Figure S5:**
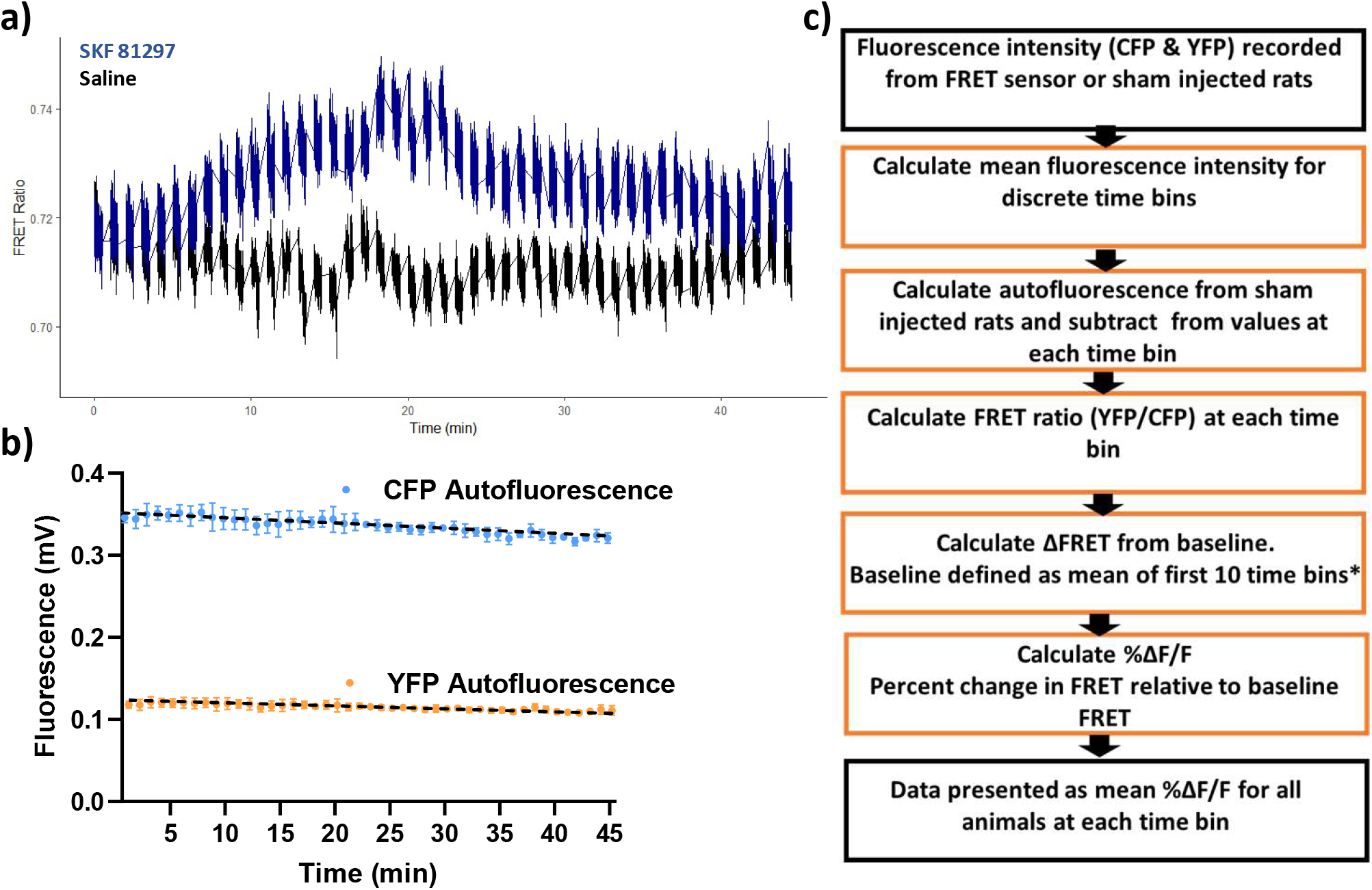
Acquisition and processing of photometry data. **(a)** FRET ratio of unprocessed fluorescent data. As described in *Methods*, data were acquired at 100 Hz in discrete 30-s intervals, with the laser turned off between sampling intervals. **(b)** Time course showing autofluorescence in CFP and YFP channels, recorded in the same way from control rats that were not injected with virus. Data displayed represents the mean ± SEM at each time (n = 2 rats). The dotted line represents a linear fit used to calculate the expected autofluorescence in each channel at a specified time point. **(c)** Flow chart describing the steps of data processing used to calculate the %ΔF/F measure.

**Figure S6:**
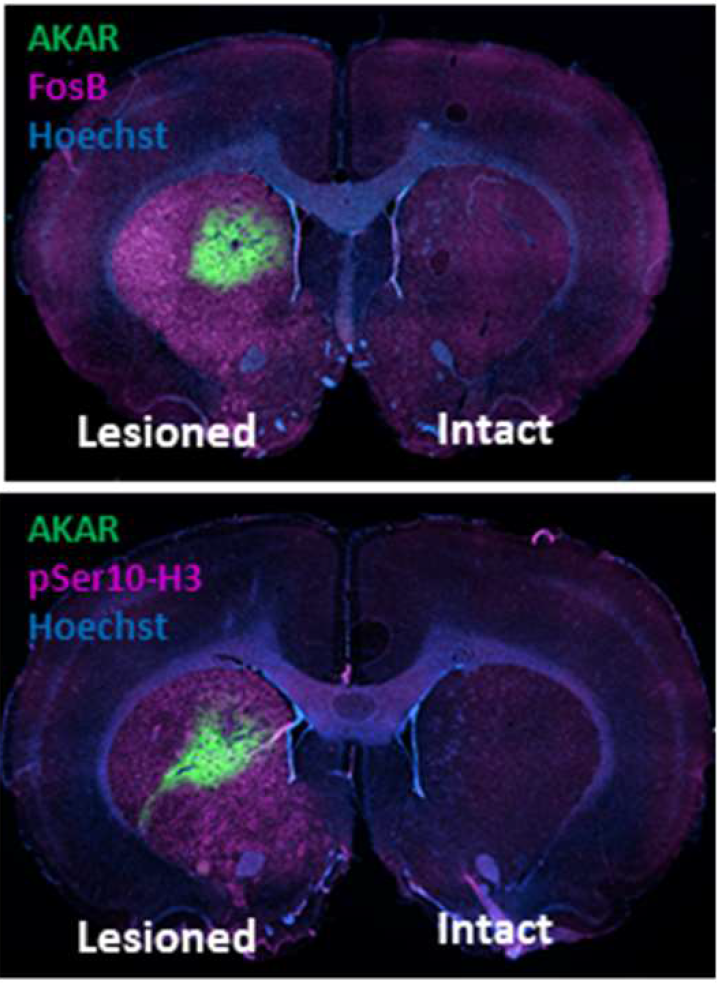
Immunofluorescent staining for markers of dyskinesia in 6-OHDA lesioned, L-DOPA treated rats. Representative images showing immunolabelling of FosB and serine-10 phosphorylated histone 3 (pSer10-H3) in tissue from 6-OHDA lesioned rats after 2 weeks of daily treatment with L-DOPA/benserazide (6/15 mg/kg s.c.) (n = 5 rats).

## References

1 Vassilatis, D. K. et al. The G protein-coupled receptor repertoires of human and mouse. Proceedings of the National Academy of Sciences of the United States of America 100, 4903–4908, doi:10.1073/pnas.0230374100 (2003).

2 Day, R. N., Tao, W. & Dunn, K. W. A simple approach for measuring FRET in fluorescent biosensors using two-photon microscopy. Nature protocols 11, 2066–2080, doi:10.1038/nprot.2016.121 (2016).

3 Goto, A. et al. Circuit-dependent striatal PKA and ERK signaling underlies rapid behavioral shift in mating reaction of male mice. Proceedings of the National Academy of Sciences of the United States of America 112, 6718–6723, doi:10.1073/pnas.1507121112 (2015).

4 Lee, S. J., Chen, Y., Lodder, B. & Sabatini, B. L. Monitoring Behaviorally Induced Biochemical Changes Using Fluorescence Lifetime Photometry. 13, doi:10.3389/fnins.2019.00766 (2019).

5 Ma, L. et al. A Highly Sensitive A-Kinase Activity Reporter for Imaging Neuromodulatory Events in Awake Mice. Neuron 99, 665–679.e665, doi:https://doi.org/10.1016/j.neuron.2018.07.020 (2018).

6 Laviv, T. et al. In Vivo Imaging of the Coupling between Neuronal and CREB Activity in the Mouse Brain. Neuron, doi:https://doi.org/10.1016/j.neuron.2019.11.028 (2019).

7 Cui, G. et al. Concurrent activation of striatal direct and indirect pathways during action initiation. Nature 494, 238–242, doi:10.1038/nature11846 (2013).

8 Gunaydin, Lisa A. et al. Natural Neural Projection Dynamics Underlying Social Behavior. Cell 157, 1535–1551, doi:https://doi.org/10.1016/j.cell.2014.05.017 (2014).

9 Muir, J. et al. In Vivo Fiber Photometry Reveals Signature of Future Stress Susceptibility in Nucleus Accumbens. Neuropsychopharmacology 43, 255–263, doi:10.1038/npp.2017.122 (2018).

10 Reed, S. J. et al. Coordinated Reductions in Excitatory Input to the Nucleus Accumbens Underlie Food Consumption. Neuron 99, 1260–1273.e1264, doi:https://doi.org/10.1016/j.neuron.2018.07.051 (2018).

11 Miyawaki, A. Development of Probes for Cellular Functions Using Fluorescent Proteins and Fluorescence Resonance Energy Transfer. 80, 357–373, doi:10.1146/annurev-biochem-072909-094736 (2011).

12 Klarenbeek, J., Goedhart, J., van Batenburg, A., Groenewald, D. & Jalink, K. Fourth-generation epac-based FRET sensors for cAMP feature exceptional brightness, photostability and dynamic range: characterization of dedicated sensors for FLIM, for ratiometry and with high affinity. PLoS One 10, e0122513, doi:10.1371/journal.pone.0122513 (2015).

13 Komatsu, N. et al. Development of an optimized backbone of FRET biosensors for kinases and GTPases. Molecular biology of the cell 22, 4647–4656, doi:10.1091/mbc.E11-01-0072 (2011).

14 Nobis, M. et al. A RhoA-FRET Biosensor Mouse for Intravital Imaging in Normal Tissue Homeostasis and Disease Contexts. Cell reports 21, 274–288, doi:10.1016/j.celrep.2017.09.022 (2017).

15 Peng, Q. et al. Coordinated histone modifications and chromatin reorganization in a single cell revealed by FRET biosensors. Proceedings of the National Academy of Sciences of the United States of America 115, E11681–e11690, doi:10.1073/pnas.1811818115 (2018).

16 Hiratsuka, T. et al. Live imaging of extracellular signal-regulated kinase and protein kinase A activities during thrombus formation in mice expressing biosensors based on Förster resonance energy transfer. Journal of thrombosis and haemostasis : JTH 15, 1487–1499, doi:10.1111/jth.13723 (2017).

17 Kamioka, Y. et al. Live imaging of protein kinase activities in transgenic mice expressing FRET biosensors. Cell structure and function 37, 65–73, doi:10.1247/csf.11045 (2012).

18 Beaulieu, J.-M. & Gainetdinov, R. R. The Physiology, Signaling, and Pharmacology of Dopamine Receptors. 63, 182–217, doi:10.1124/pr.110.002642 % (2011).

19 Nishi, A., Kuroiwa, M. & Shuto, T. Mechanisms for the Modulation of Dopamine D1 Receptor Signaling in Striatal Neurons. 5, doi:10.3389/fnana.2011.00043 (2011).

20 Price, C. J., Kim, P. & Raymond, L. A. D1 Dopamine Receptor-Induced Cyclic AMP-Dependent Protein Kinase Phosphorylation and Potentiation of Striatal Glutamate Receptors. 73, 2441–2446, doi:10.1046/j.1471-4159.1999.0732441.x (1999).

21 Willuhn, I. & Steiner, H. Motor-skill learning in a novel running-wheel task is dependent on D1 dopamine receptors in the striatum. Neuroscience 153, 249–258, doi:https://doi.org/10.1016/j.neuroscience.2008.01.041 (2008).

22 Alcacer, C. et al. Gα (olf) mutation allows parsing the role of cAMP-dependent and extracellular signal-regulated kinase-dependent signaling in L-3,4-dihydroxyphenylalanine-induced dyskinesia. The Journal of neuroscience : the official journal of the Society for Neuroscience 32, 5900–5910, doi:10.1523/jneurosci.0837-12.2012 (2012).

23 Corvol, J. C. et al. Persistent increase in olfactory type G-protein alpha subunit levels may underlie D1 receptor functional hypersensitivity in Parkinson disease. The Journal of neuroscience : the official journal of the Society for Neuroscience 24, 7007–7014, doi:10.1523/jneurosci.0676-04.2004 (2004).

24 Gerfen, C. R., Miyachi, S., Paletzki, R. & Brown, P. D1 dopamine receptor supersensitivity in the dopamine-depleted striatum results from a switch in the regulation of ERK1/2/MAP kinase. The Journal of neuroscience : the official journal of the Society for Neuroscience 22, 5042–5054 (2002).

25 Morigaki, R., Okita, S. & Goto, S. Dopamine-Induced Changes in Gαolf Protein Levels in Striatonigral and Striatopallidal Medium Spiny Neurons Underlie the Genesis of l-DOPA-Induced Dyskinesia in Parkinsonian Mice. Frontiers in cellular neuroscience 11, 26, doi:10.3389/fncel.2017.00026 (2017).

26 Darmopil, S., Martin, A. B., De Diego, I. R., Ares, S. & Moratalla, R. Genetic inactivation of dopamine D1 but not D2 receptors inhibits L-DOPA-induced dyskinesia and histone activation. Biological psychiatry 66, 603–613, doi:10.1016/j.biopsych.2009.04.025 (2009).

27 Lebel, M., Chagniel, L., Bureau, G. & Cyr, M. Striatal inhibition of PKA prevents levodopa-induced behavioural and molecular changes in the hemiparkinsonian rat. Neurobiology of disease 38, 59–67, doi:10.1016/j.nbd.2009.12.027 (2010).

28 Santini, E. et al. Critical Involvement of cAMP/DARPP-32 and Extracellular Signal-Regulated Protein Kinase Signaling in l-DOPA-Induced Dyskinesia. 27, 6995–7005, doi:10.1523/JNEUROSCI.0852-07.2007 %J The Journal of Neuroscience (2007).

29 Muntean, B. S. et al. Interrogating the Spatiotemporal Landscape of Neuromodulatory GPCR Signaling by Real-Time Imaging of cAMP in Intact Neurons and Circuits. Cell reports 22, 255–268, doi:10.1016/j.celrep.2017.12.022 (2018).

30 Graveland, G. A. & Difiglia, M. The frequency and distribution of medium-sized neurons with indented nuclei in the primate and rodent neostriatum. Brain Research 327, 307–311, doi:https://doi.org/10.1016/0006-8993(85)91524-0 (1985).

31 Ouimet, C. C., Langley-Gullion, K. C. & Greengard, P. Quantitative immunocytochemistry of DARPP-32-expressing neurons in the rat caudatoputamen. Brain Research 808, 8–12, doi:https://doi.org/10.1016/S0006-8993(98)00724-0 (1998).

32 Langley, K. C., Bergson, C., Greengard, P. & Ouimet, C. C. Co-localization of the D1 dopamine receptor in a subset of DARPP-32-containing neurons in rat caudate–putamen. Neuroscience 78, 977–983, doi:https://doi.org/10.1016/S0306-4522(96)00583-0 (1997).

33 Matamales, M. et al. Striatal Medium-Sized Spiny Neurons: Identification by Nuclear Staining and Study of Neuronal Subpopulations in BAC Transgenic Mice. PLoS One 4, e4770, doi:10.1371/journal.pone.0004770 (2009).

34 Yapo, C. et al. Detection of phasic dopamine by D1 and D2 striatal medium spiny neurons. The Journal of physiology 595, 7451–7475, doi:10.1113/jp274475 (2017).

35 Poewe, W. et al. Parkinson disease. Nature Reviews Disease Primers 3, 17013, doi:10.1038/nrdp.2017.13 (2017).

36 Cheng, H. C., Ulane, C. M. & Burke, R. E. Clinical progression in Parkinson disease and the neurobiology of axons. Annals of neurology 67, 715–725, doi:10.1002/ana.21995 (2010).

37 Delfino, M. A. et al. Behavioral sensitization to different dopamine agonists in a parkinsonian rodent model of drug-induced dyskinesias. Behavioural Brain Research 152, 297–306, doi:https://doi.org/10.1016/j.bbr.2003.10.009 (2004).

38 Ungerstedt, U. 6-hydroxy-dopamine induced degeneration of central monoamine neurons. European Journal of Pharmacology 5, 107–110, doi:https://doi.org/10.1016/0014-2999(68)90164-7 (1968).

39 Huot, P., Johnston, T. H., Koprich, J. B., Fox, S. H. & Brotchie, J. M. The Pharmacology of L-DOPA-Induced Dyskinesia in Parkinson’s Disease. 65, 171–222, doi:10.1124/pr.111.005678 %J Pharmacological Reviews (2013).

40 Jenner, P. Molecular mechanisms of L-DOPA-induced dyskinesia. Nature Reviews Neuroscience 9, 665–677, doi:10.1038/nrn2471 (2008).

41 Girasole, A. E. et al. A Subpopulation of Striatal Neurons Mediates Levodopa-Induced Dyskinesia. Neuron 97, 787–795.e786, doi:https://doi.org/10.1016/j.neuron.2018.01.017 (2018).

42 Keifman, E. et al. Optostimulation of striatonigral terminals in substantia nigra induces dyskinesia that increases after L-DOPA in a mouse model of Parkinson’s disease. 176, 2146–2161, doi:10.1111/bph.14663 (2019).

43 Ryan, M. B., Bair-Marshall, C. & Nelson, A. B. Aberrant Striatal Activity in Parkinsonism and Levodopa-Induced Dyskinesia. Cell reports 23, 3438–3446.e3435, doi:10.1016/j.celrep.2018.05.059 (2018).

44 Conti, M. M. et al. Effect of tricyclic antidepressants on L-DOPA-induced dyskinesia and motor improvement in hemi-parkinsonian rats. Pharmacology, biochemistry, and behavior 142, 64–71, doi:10.1016/j.pbb.2016.01.004 (2016).

45 Frouni, I. et al. Effect of the selective 5-HT2A receptor antagonist EMD-281,014 on L-DOPA-induced abnormal involuntary movements in the 6-OHDA-lesioned rat. Experimental brain research 237, 29–36, doi:10.1007/s00221-018-5390-4 (2019).

46 Cenci, M. A. & Lundblad, M. Ratings of L-DOPA-induced dyskinesia in the unilateral 6-OHDA lesion model of Parkinson's disease in rats and mice. Current protocols in neuroscience Chapter 9, Unit 9.25, doi:10.1002/0471142301.ns0925s41 (2007).

47 Feyder, M. et al. A Role for Mitogen- and Stress-Activated Kinase 1 in L-DOPA–Induced Dyskinesia and ΔFosB Expression. Biological psychiatry 79, 362–371, doi:https://doi.org/10.1016/j.biopsych.2014.07.019 (2016).

48 Pavón, N., Martín, A. B., Mendialdua, A. & Moratalla, R. ERK Phosphorylation and FosB Expression Are Associated with L-DOPA-Induced Dyskinesia in Hemiparkinsonian Mice. Biological psychiatry 59, 64–74, doi:https://doi.org/10.1016/j.biopsych.2005.05.044 (2006).

49 Santini, E. et al. l-DOPA activates ERK signaling and phosphorylates histone H3 in the striatonigral medium spiny neurons of hemiparkinsonian mice. 108, 621–633, doi:10.1111/j.1471-4159.2008.05831.x (2009).

50 Feyder, M., Bonito Oliva, A. & Fisone, G. L-DOPA-Induced Dyskinesia and Abnormal Signaling in Striatal Medium Spiny Neurons: Focus on Dopamine D1 Receptor-Mediated Transmission. 5, doi:10.3389/fnbeh.2011.00071 (2011).

51 Spigolon, G. & Fisone, G. J. J. o. N. T. Signal transduction in l-DOPA-induced dyskinesia: from receptor sensitization to abnormal gene expression. 125, 1171–1186, doi:10.1007/s00702-018-1847-7 (2018).

52 Mariani, L.-L., Longueville, S., Girault, J.-A., Hervé, D. & Gervasi, N. Differential enhancement of ERK, PKA and Ca2+ signaling in direct and indirect striatal neurons of Parkinsonian mice. Neurobiology of disease 130, 104506, doi:https://doi.org/10.1016/j.nbd.2019.104506 (2019).

53 Akemann, W. et al. Imaging neural circuit dynamics with a voltage-sensitive fluorescent protein. 108, 2323–2337, doi:10.1152/jn.00452.2012 (2012).

54 Kamioka, Y. et al. Live Imaging of Protein Kinase Activities in Transgenic Mice Expressing FRET Biosensors. Cell structure and function 37, 65–73, doi:10.1247/csf.11045 (2012).

55 Tao, W. et al. A practical method for monitoring FRET-based biosensors in living animals using two-photon microscopy. American journal of physiology. Cell physiology 309, C724–735, doi:10.1152/ajpcell.00182.2015 (2015).

56 Guo, Q. et al. Multi-channel fiber photometry for population neuronal activity recording. Biomed Opt Express 6, 3919–3931, doi:10.1364/BOE.6.003919 (2015).

57 Kim, C. K. et al. Simultaneous fast measurement of circuit dynamics at multiple sites across the mammalian brain. Nature Methods 13, 325–328, doi:10.1038/nmeth.3770 (2016).

58 Cavero, I., Massingham, R. & Lefevere-Borg, F. Peripheral dopamine receptors, potential targets for a new class of antihypertensive agents. Part II: Sites and mechanisms of action of dopamine receptor agonists. Life sciences 31, 1059–1069, doi:10.1016/0024-3205(82)90078-9 (1982).

59 Cavero, I., Thiry, C., Pratz, J. & Lawson, K. Cardiovascular characterization of DA-1 and DA-2 dopamine receptor agonists in anesthetized rats. Clinical and experimental hypertension. Part A, Theory and practice 9, 931–952, doi:10.3109/10641968709161458 (1987).

60 Van der Niepen, P., Dupont, A. G., Finne, E. & Six, R. O. Hypotensive and regional haemodynamic effects of fenoldopam and quinpirole in the anaesthetized rat. Journal of hypertension. Supplement : official journal of the International Society of Hypertension 6, S687–689, doi:10.1097/00004872-198812040-00216 (1988).

61 Ma, Y. et al. Wide-field optical mapping of neural activity and brain haemodynamics: considerations and novel approaches. Philosophical transactions of the Royal Society of London. Series B, Biological sciences 371, doi:10.1098/rstb.2015.0360 (2016).

62 Valley, M. T. et al. Separation of hemodynamic signals from GCaMP fluorescence measured with wide-field imaging. Journal of neurophysiology 123, 356–366, doi:10.1152/jn.00304.2019 (2020).

63 Yasuda, R. in Neurophotonics and Biomedical Spectroscopy (eds Robert R. Alfano & Lingyan Shi) 53–64 (Elsevier, 2019).

64 Lee, S. J. et al. Cell-type specific asynchronous modulation of PKA by dopamine during reward based learning. 839035, doi:10.1101/839035 %J bioRxiv (2019).

65 Aubert, I. et al. Increased D1 dopamine receptor signaling in levodopa-induced dyskinesia. 57, 17–26, doi:10.1002/ana.20296 (2005).

66 Corvol, J.-C. et al. Persistent Increase in Olfactory Type G-Protein α Subunit Levels May Underlie D1 Receptor Functional Hypersensitivity in Parkinson Disease. 24, 7007–7014, doi:10.1523/JNEUROSCI.0676-04.2004 % (2004).

67 Guigoni, C., Doudnikoff, E., Li, Q., Bloch, B. & Bezard, E. Altered D1 dopamine receptor trafficking in parkinsonian and dyskinetic non-human primates. Neurobiology of disease 26, 452–463, doi:https://doi.org/10.1016/j.nbd.2007.02.001 (2007).

68 Rangel-Barajas, C. et al. L-DOPA-induced dyskinesia in hemiparkinsonian rats is associated with up-regulation of adenylyl cyclase type V/VI and increased GABA release in the substantia nigra reticulata. Neurobiology of disease 41, 51–61, doi:10.1016/j.nbd.2010.08.018 (2011).

69 Chen, G. et al. Antidyskinetic Effects of MEK Inhibitor Are Associated with Multiple Neurochemical Alterations in the Striatum of Hemiparkinsonian Rats. Frontiers in neuroscience 11, 112, doi:10.3389/fnins.2017.00112 (2017).

70 Fieblinger, T. et al. Cell type-specific plasticity of striatal projection neurons in parkinsonism and L-DOPA-induced dyskinesia. Nature Communications 5, 5316, doi:10.1038/ncomms6316 (2014).

71 Calabresi, P. et al. Dopamine and cAMP-regulated phosphoprotein 32 kDa controls both striatal long-term depression and long-term potentiation, opposing forms of synaptic plasticity. The Journal of neuroscience : the official journal of the Society for Neuroscience 20, 8443–8451 (2000).

72 Picconi, B. et al. Loss of bidirectional striatal synaptic plasticity in L-DOPA-induced dyskinesia. Nature neuroscience 6, 501–506, doi:10.1038/nn1040 (2003).

73 Picconi, B. et al. l-DOPA dosage is critically involved in dyskinesia via loss of synaptic depotentiation. Neurobiology of disease 29, 327–335, doi:10.1016/j.nbd.2007.10.001 (2008).

74 Berton, O. et al. Striatal overexpression of ΔJunD resets L-DOPA-induced dyskinesia in a primate model of Parkinson disease. Biological psychiatry 66, 554–561, doi:10.1016/j.biopsych.2009.04.005 (2009).

75 Cao, X. et al. Striatal overexpression of ΔFosB reproduces chronic levodopa-induced involuntary movements. The Journal of neuroscience : the official journal of the Society for Neuroscience 30, 7335–7343, doi:10.1523/jneurosci.0252-10.2010 (2010).

76 Beck, G. et al. Role of striatal ΔFosB in l-Dopa-induced dyskinesias of parkinsonian nonhuman primates. Proceedings of the National Academy of Sciences of the United States of America 116, 18664–18672, doi:10.1073/pnas.1907810116 (2019).

77 Sohn, J. et al. A Single Vector Platform for High-Level Gene Transduction of Central Neurons: Adeno-Associated Virus Vector Equipped with the Tet-Off System. PLoS One 12, e0169611–e0169611, doi:10.1371/journal.pone.0169611 (2017).

78 Burger, C. & Nash, K. R. in Gene Therapy for Neurological Disorders: Methods and Protocols (ed Fredric P. Manfredsson) 95–106 (Springer New York, 2016).

79 Brewer, G. J. & Torricelli, J. R. Isolation and culture of adult neurons and neurospheres. Nature protocols 2, 1490–1498, doi:10.1038/nprot.2007.207 (2007).

80 Bardy, C. et al. Neuronal medium that supports basic synaptic functions and activity of human neurons in vitro. Proceedings of the National Academy of Sciences of the United States of America 112, E2725–E2734, doi:10.1073/pnas.1504393112 (2015).

81 Schallert, T., Fleming, S. M., Leasure, J. L., Tillerson, J. L. & Bland, S. T. CNS plasticity and assessment of forelimb sensorimotor outcome in unilateral rat models of stroke, cortical ablation, parkinsonism and spinal cord injury. Neuropharmacology 39, 777–787, doi:10.1016/s0028-3908(00)00005-8 (2000).

